# Fluoride triggers lysis in *Streptococcus mutans* by inhibition of Clp protease complex leading to an unabated competence cascade

**DOI:** 10.1101/2025.02.28.640611

**Authors:** Aditya Banerjee, Sophie Pickelner, Faith Smith, Livia M.A. Tenuta, Randy B. Stockbridge

## Abstract

Fluoride has long been known to possess antimicrobial properties, triggering extensive lysis in diverse bacterial species. Surprisingly, given the ubiquity of fluoride in oral healthcare, the underlying mechanisms by which fluoride kills bacteria remain unknown. Using dental pathogen *Streptococcus mutans* as a model, we show that fluoride elicits an uncontrolled stress response characterized by upregulation of competence pathways and extensive cell wall degradation. While limited autolysis is adaptive, we show that fluoride subverts the typical tight temporal control of the competence-associated alternative sigma factor ComX by specifically inhibiting assembly and activity of Clp ATPases, thus disrupting proteolytic degradation of ComX. Phenylalanine partially restores Clp protease activity, explaining the frequent presence of a gene encoding chorismate mutase in fluoride-responsive operons. Together, these findings reveal the molecular mechanism that underlie fluoride-dependent lysis in bacteria, and identify new pathways that could be exploited to potentiate the antimicrobial effects of oral fluoride.

## Introduction

Fluoride is ubiquitous in the microbial biosphere, and most bacteria evolved in the presence of physiologically relevant amounts of this inhibitory ion (∼50 μM - 1 mM)^1^. Human activity has vastly increased the incidence of fluoride in certain environments, including the nearly universal use of fluoride in oral healthcare, and the ubiquitous use of fluorinated organic compounds in medicine, commerce, and industry^2^. Following this proliferation, in recent years, interest in exploiting microbial fluoride sensitivity or resistance mechanisms has burgeoned across multiple different fields. For example, in dentistry, reducing the fluoride concentrations used in oral healthcare has been an important goal, and methods to potentiate the antibacterial effect of oral fluoride are being sought^3–6^. Meanwhile, synthetic biologists seek to engineer microbes to synthesize novel fluorinated compounds, or to degrade organofluorine pollutants^7–11^. Both degradation and synthetic pathways involve fluoride ion, and microbial tolerance to cytosolic fluoride presents a key challenge to both^2, 9, 12^. In light of these diverse emerging applications, a molecular understanding of specific mechanisms of bacterial resistance and susceptibility to fluoride is becoming increasingly important.

The molecular components of the fluoride stress response vary among bacteria. For most microbes, the first lines of defense against fluoride are membrane-embedded fluoride exporters, including the fluoride/proton antiporter CLC^F^, or fluoride channels from the Fluc family^13^. These proteins extrude cytosolic fluoride by harnessing the proton gradient and the positive-outside membrane potential, respectively, to maintain fluoride at sub-inhibitory levels^14, 15^. In addition to expressing export proteins, many bacteria resist fluoride by synthesizing additional copies of enzymes that are inhibited by fluoride, such as enolase, pyrophosphatase, and ATPases^2, 16–18^. Fluoride also perturbs pH homeostasis, metal ion homeostasis, and glycolytic metabolism^9, 19, 20^. Accordingly, various microbes increase expression of proteins like the Na^+^/H^+^ antiporter to regulate intracellular pH, metal ion transporters, and proteins involved in non-oxidative metabolism to compensate^16, 17, 21^. In some bacteria, these cellular responses are orchestrated by fluoride riboswitches, but in others, the regulatory mechanisms remain unknown^16^.

For many bacteria, these responses are sufficient to endure the fluoride challenge. Fluoride inhibition of key enzymes, including those involved in glycolysis and phosphoryl group transfer, is reversible^22–24^; thus, although fluoride exerts a bacteriostatic effect on microbes such as *Escherichia coli* and *Pseudomonas putida*, cellular growth resumes once cytosolic fluoride drops below the inhibitory constants of these enzymes^7, 9, 19^. For other species, however, fluoride is bactericidal^4, 25–27^. This phenomenon was first observed in the 1970’s, when Marquis and colleagues reported that fluoride triggered the “massive lysis” of several microbial species, including *Bacillus subtilis*, *Neisseria subflava,* and oral pathogens *Streptococcus mutans* and *Streptococcus sanguinis,* in a process that involved degradation of cell wall peptidoglycan^25^. For all of these strains, autolysis of a sub-population is an adaptive stress response that provides nutrients and genetic material to sister cells^28^. However, fluoride stood out as instigating a more dramatic, population-level lysis event. Our recent studies underscore the dramatic sensitivity of *S. mutans* to intracellular fluoride accumulation. For cells with a genetic knockout of the fluoride exporter CLC^F^ (ΔCLC^F^), the entire population rapidly lyses in the presence of one millimolar fluoride, with visible and almost complete degradation of the cell walls^4^. However, in contrast to the well-established mechanisms for temporary growth inhibition by fluoride^20, 29^, the fundamental mechanisms responsible for fluoride-dependent killing of bacteria have never been determined.

In this work, we uncover the molecular basis of fluoride’s bactericidal effect for the first time. By exploiting a fluoride export-deficient ΔCLC^F^ mutant of *S. mutans*, we show that fluoride, like other sources of cellular stress, triggers a stress response including the upregulation of competence-associated cell wall hydrolases via the competence-related alternative sigma factor ComX. However, unlike other cellular stressors, fluoride subverts cellular mechanisms to turn *off* these competence pathways by directly inhibiting the assembly of the Clp protease complex that degrades ComX. Finally, we show that phenylalanine and tyrosine, but not tryptophan, partially rescue ClpXP-mediated proteolysis of ComX, addressing a long-standing question about the association between chorismate mutase and fluoride resistance, and establishing aromatic amino acid synthesis as a novel fluoride resistance mechanism in bacteria.

## Results

### Fluoride stress triggers cell lysis through the misregulation of competence programs

We have previously shown that a fluoride export-deficient mutant of *S. mutans* is killed by 0.3 mM NaF^4^. This strain bears deletions of *smu_1289* and *smu_1290,* which encode the fluoride exporter CLC^F^, and we refer to this strain as ΔCLC^F^. We confirmed that fluoride triggers similar population-level cell death in WT *S. mutans*, albeit at a higher fluoride concentration (**Supplementary Figure 1**). WT strains of other oral *Bacilliota,* including diverse *Streptococci* and *Lactococcus salivarius*, also exhibited extensive cell death like *S. mutans* upon fluoride challenge (**Supplementary Figure 2**).

We elected to use the ΔCLC^F^ *S. mutans* strain as a responsive model to investigate the cellular mechanisms of fluoride-induced bacterial death. We first sought to understand whether fluoride’s bactericidal effect occurs due to cell lysis, and whether this differs from other lethal stressors. We calibrated three different stressors, heat, chloramphenicol, and fluoride, to induce comparable levels of cell death in *S. mutans*, assessed by replating on recovery plates after the challenge and counting colony forming units 48 hours later (**Supplementary Figure 3**). Visualization of cell morphology after treatment using transmission electron microscopy (TEM, **Figure 1a**) shows only a few cells with compromised cell walls after heat or chloramphenicol treatments. The sample treated with 0.3 mM NaF, however, shows far more cell wall damage, including many cells for which the cell wall has been entirely degraded. These results suggest that, among these diverse stressors, fluoride is unique in the extent of cell wall degradation. As expected from the visual lysis phenotype, we observed a significant increase in the expression of numerous autolysin-encoding genes in ΔCLC^F^ *S. mutans* treated with 0.3 mM NaF, including muramidases (*smu_704c*, *smu_76)*, endolysin (*smu_707*), lytic transglycosylase (*smu_2147*) and hydrolases (*smu_1070c*, *smu_574*, *smu_1700*) (**Figure 1b**). At the same time, autolysis attenuators^30, 31^ *lytR* and *dltD* did not exhibit an increase in expression. These results confirm the heightened activity of lytic pathways in the fluoride-treated cells.

**Figure 1.**
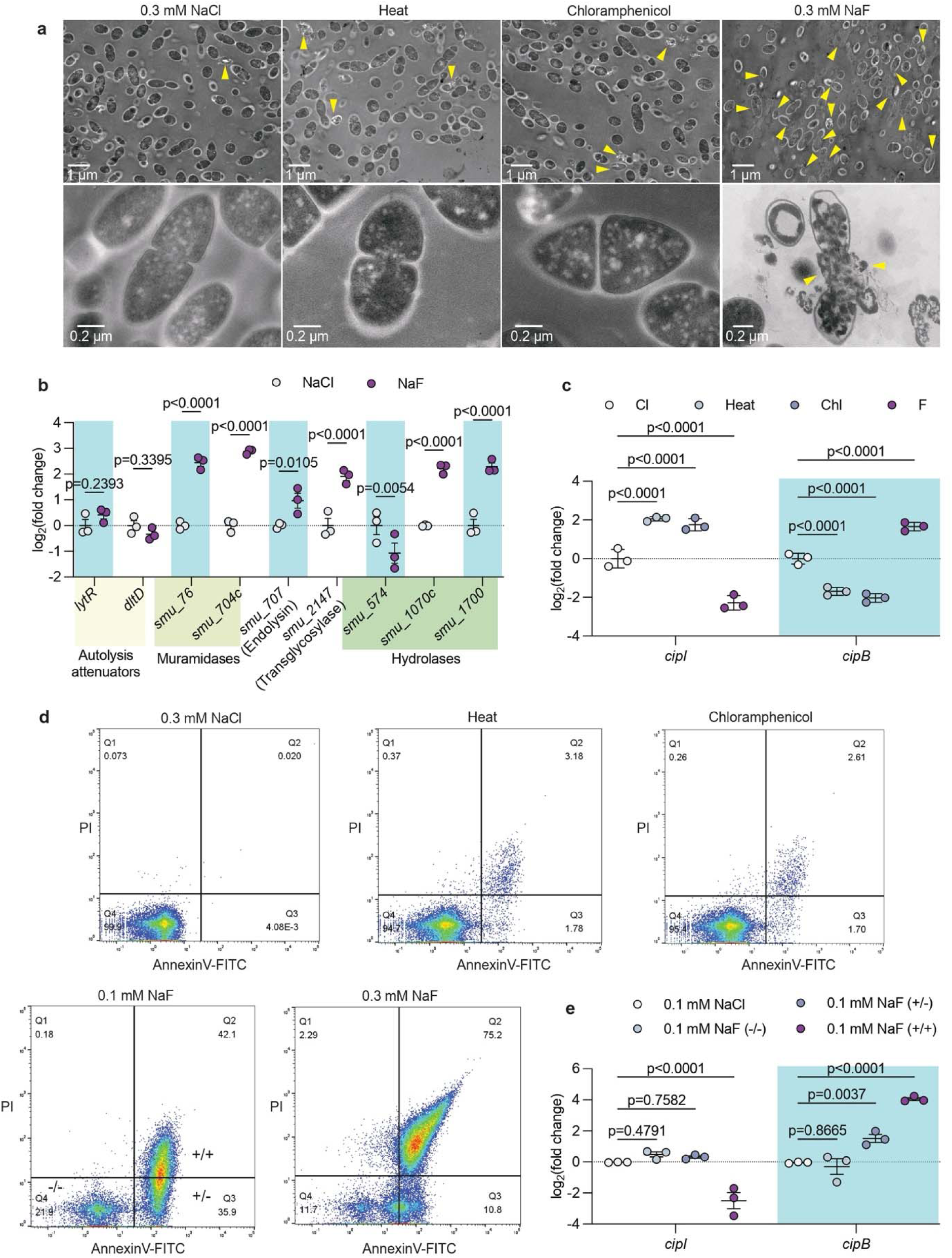
Fluoride stress triggers cell lysis through misregulation of competence programs. For all experiments, the ΔCLC^F^ *S. mutans* strain was treated with 0.3 mM NaCl (control), 0.3 mM NaF, chloramphenicol, or heat, with dosages calibrated as described in the *Methods* and **Supplementary Figure 2**. **a.** TEM images of cells after treatment with NaCl, NaF, chloramphenicol or heat. Yellow arrows indicate cell wall disruptions or dispersed cytosolic material. **b.** Expression of selected autolysis-associated genes in control and fluoride-treated cells. Datapoints represent mean and SEM of three independent biological replicates (*n*=3). Significance is calculated using two-way analysis of variance (ANOVA) followed by Fisher’s LSD test. Gene expression is normalized relative to housekeeping gene *gyrA* and the chloride control sample. **c**. Expression of *cipI* and *cipB* in control, fluoride-, heat-, and chloramphenicol-treated cells, with normalization and statistics as in panel **b**. **d.** Sorted cell populations for control cells and cells after treatment with 0.1 or 0.3 mM NaF, heat, or chloramphenicol, stained with AnnexinV-FITC and propidium iodide (PI). **e.** Expression of *cipI* and *cipB* in the three sorted populations, -/-, +/- and +/+, after 0.1 mM NaF treatment, with normalization and statistics as in panel **b.**

The pathways that govern stress-induced autolysis in *S. mutans* under stress have been determined^32–34^. Briefly, stress is sensed by two-component signal transduction systems at the membrane, of which LiaFSR and ComDE are the most important. The ComDE pathway ultimately produces the competence-associated bacteriocins, including CipB^35–38^. Meanwhile, induction of the LiaFSR pathway produces CipB’s immunity peptide, CipI, which sequesters CipB as part of the feedback mechanism to limit autolysis to a subpopulation of cells^39^. After heat and chloramphenicol stress, we observed induction of the immunity peptide *cipI* and a decrease in *cipB*, reflecting the activation of feedback mechanisms to curb competence pathways (**Figure 1c**). In contrast, under fluoride stress, the expression pattern was reversed, with significantly higher *cipB* expression and significantly reduced *cipI* expression. This suggests that CipI/CipB imbalance is a signature of the misregulated lysis that occurs after fluoride treatment.

To further establish a link between *cipI* and *cipB* expression and fluoride-induced cell death, we exposed *S. mutans* to sublethal 0.1 mM NaF, and stained the cells with AnnexinV-FITC and propidium iodide (PI). Normally excluded by the cell wall, AnnexinV-FITC is only able to access and label phosphatidylserine of the inner membrane when the cell wall is leaky, consistent with competence or cell wall remodeling/degradation. PI is normally excluded by the plasma membrane, and PI staining therefore reflects membrane damage. We then used cell sorting to isolate three populations of cells: cells with intact cell walls and plasma membrane (-/-), moribund cells with permeable cell walls and damaged membranes (+/+), and an intermediate population of cells that maintain an impermeable membrane, but have damaged cell walls (+/-). The -/- and +/- populations were able to re-grow on recovery plates, whereas cells from the +/+ fraction could not (**Supplementary Figure 4**). Relative to the other two populations, cells from the +/+ fraction exhibited elevated levels of organic acids and bactericidal metabolites, and lower levels of glutathione (**Supplementary Figure 5)**, all of which are associated with cellular lysis and stress^40–44^.

Cells exposed to 0.1 mM NaF were primarily distributed across the AnnexinV-positive (+/-, ∼36%) and the PI/AnnexinV positive quadrants (+/+, ∼42%), indicating widespread cell wall permeability in this population (**Figure 1d**). Increasing the fluoride dose to 0.3 mM NaF increased the extent of the cell wall and membrane damage, so that the majority (∼75%) of cells were positive for both PI and AnnexinV. In contrast, for cells treated with heat or chloramphenicol, the largest proportion occupied the double negative quadrant, but were nonetheless inviable (**Supplementary Figure 4**). Thus, the extensive membrane and cell wall damage of the fluoride treated population is not a general feature of *S. mutans* cell death, consistent with previous observations^45^ (**Figure 1d**). We evaluated the three sorted cell populations from the sample treated with 0.1 mM NaF for *cipI* and *cipB* expression, which confirmed that the differential expression of these genes is correlated with the wall and membrane phenotype (**Figure 1e**).

### The competence related sigma factor ComX mediates the fluoride stress response

To understand the origin of the CipI/CipB imbalance under fluoride stress, we next aimed to identify the mechanisms responsible for their regulation. We performed a pull-down assay of the sorted sub-populations using the *cipI* and *cipB* promoters as bait (**Figure 2a**). For the -/- and +/- populations, the primary sigma factor Sig42 was bound to the *cipI* promoter, but this was not observed in the +/+ population. At the same time, the competence-associated sigma factor ComX was associated with the CipB promoter in the +/- and +/+ populations, but was not observed in the -/- population. This transition from Sig42-mediated expression of the immunity peptide CipI to ComX-mediated expression of the bacteriocin CipB is consistent with the shift in cellular regulation towards competence.

**Figure 2.**
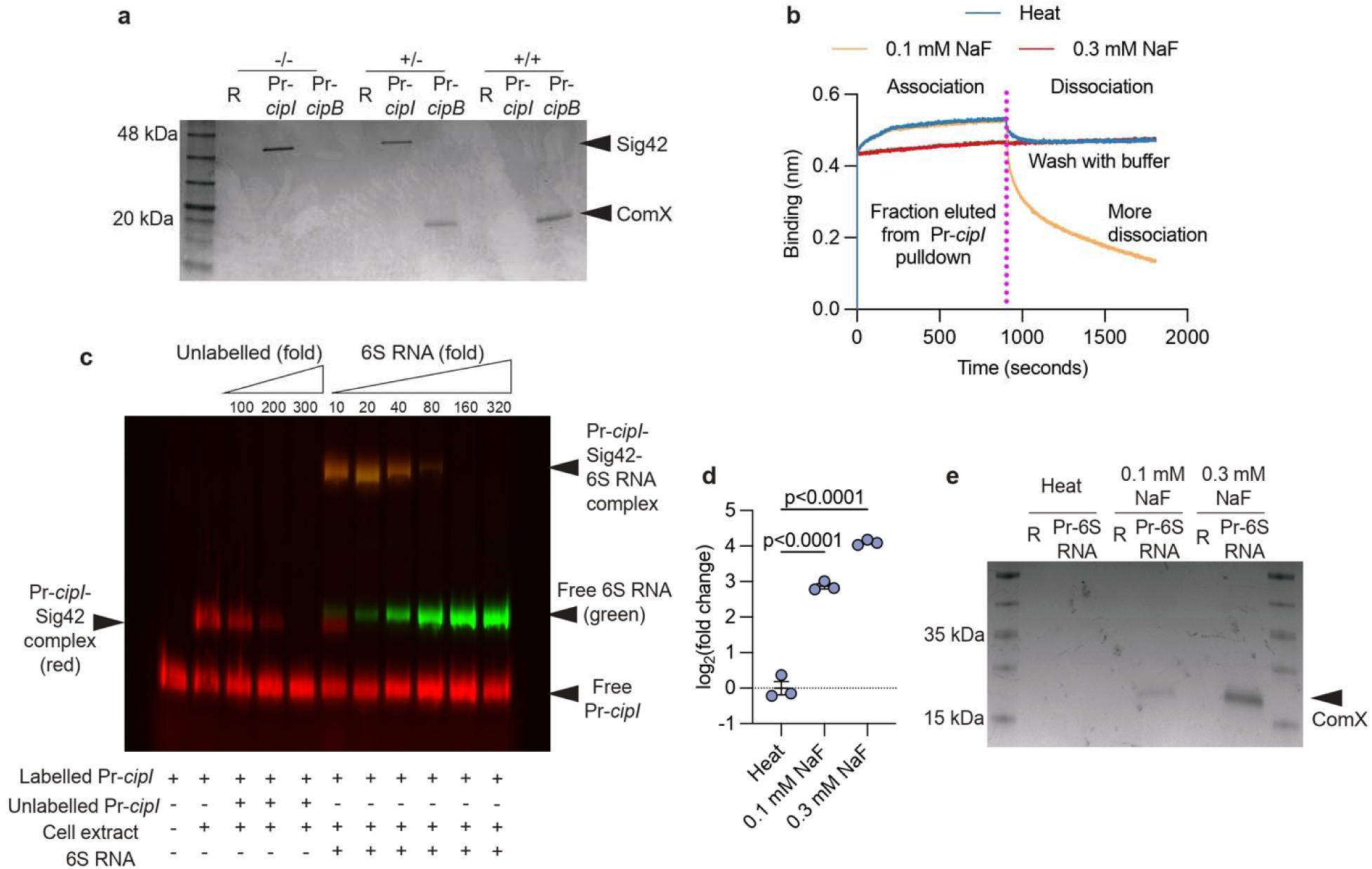
The alternative sigma factor ComX coordinates the fluoride response. **a.** SDS- PAGE after pulldown with the biotinylated promoters of *cipI* (Pr-*cipI*) and *cipB* (Pr-*cipB*), using lysates prepared from the -/-, +/- and +/+ populations from **Figure 1d**. For the lane indicated “R,” a random DNA sequence was used as bait. Bands were excised from the gel and Sig42 and ComX verified using mass spectrometry. **b.** Biolayer interferometry profile showing binding and dissociation of cellular factors to Pr*cipI* after the indicated treatments. For the binding phase, sensors bound with Pr*cipI* were incubated with the eluted fractions from the promoter pulldown assay in panel **a**. Sensors were transferred to buffer for the dissociation phase. **c.** Electrophoretic mobility shift assay (EMSA). Lane 1: Cy5-labelled Pr-*cipI*. Lane 2: Cy5-labelled Pr-*cipI* incubated with the Sig42-containing pull-down fractions from cell lysates of heat-treated cells. The mass shift is consistent with formation of the Sig42/Pr*cipI* complex. Lane 3-5: As a control, increasing unlabeled Pr-*cipI* is added to compete with the Cy5-labelled Pr-*cipI*. Lane 6-11: Sig42/Pr*cipI* complex with addition of increasing FITC-labelled 6S RNA. The molecular weight of the super-shifted band is consistent with a Pr-*cipI*-Sig42-6S RNA ternary complex. **d.** Expression of 6S RNA after heat or NaF treatment as indicated. Gene expression is normalized relative to housekeeping gene *gyrA* and the heat-treated reference sample. Datapoints represent mean and SEM of three independent biological replicates (*n*=3). Significance is calculated using one-way analysis of variance (ANOVA) followed by Fisher’s LSD test. **e.** SDS-PAGE after pulldown with the biotinylated promoter of 6S RNA (Pr-6S RNA) from lysates of cells treated with heat or NaF as indicated. For the lane indicated “R” a random DNA sequence was used as bait.

We then performed biolayer interferometry, a biophysical measurement of macromolecular complex formation/dissociation, to evaluate transcription factor dissociation from the promoter DNA sequences. The promoter sequences were incubated with the eluted fractions from the promoter pulldown assays, loading the oligonucleotides with their respective sigma factors. Subsequent incubation in buffer allows dissociation of any bound species. For cells treated with 0.1 mM NaF, we observed an enhanced dissociation signal from the *cipI* promoter, suggesting dissociation of a larger macromolecular complex (**Figure 2b**). The enhanced dissociation from the promoter was not observed under heat stress or under any conditions with the *cipB* promoter (**Figure 2b, Supplementary Figure 6**). Subsequent analysis of the *cipI* promoter pulldown fraction from the lysate of cells exposed to 0.1 mM NaF revealed an additional nucleic acid component, and RNAse and DNAse treatment identified it as an RNA (**Supplementary Figure 7**). We isolated and identified this factor as the non-coding 6S RNA, which has been implicated in switching from the primary sigma factor to alternative sigma factors during stress in *E. coli* and *B. subtilis*^46–48^. A gel mobility shift assay confirms the formation of a super-shifted complex comprising Pr-*cipI*, Sig42 and 6S RNA, which remains intact until a threshold abundance of 6S RNA, beyond which the entire complex dissociates (**Figure 2c**). These observations are all consistent with a role for 6S in mediating the switch from Sig42 to ComX regulation of *cipI*. Expression of 6S RNA is ∼2.5-fold and ∼4.0-fold higher under 0.1 and 0.3 mM NaF compared to heat (**Figure 2d**), and pull-down using the 6S RNA promoter as bait again showed ComX bound (**Figure 2e**). Thus, ComX not only induces the bacteriocin *cipB*; it also suppresses its immunity peptide *cipI* via the induction of the 6S RNA, which causes dissociation of Sig42 from the *cipI* promoter. ComX therefore plays a major role in the CipI/CipB imbalance, and related autolysis that occurs during fluoride stress.

### Fluoride inhibits Clp protease-dependent degradation of ComX

In *Streptococci*, ComX is transiently expressed during stress, triggering competence in a sub-population of cells^49^. Its expression typically peaks within ∼15 minutes, followed by rapid decay, thus limiting the timeframe for expression of competence-associated genes^49, 50^. To compare the temporal pattern of ComX expression under heat and fluoride stress, we quantified its abundance over time using targeted mass spectrometry. As anticipated, when cells are exposed to heat stress, ComX abundance peaks at 15 minutes, and returns to within 2-fold of its pre-treatment level by 30 minutes. Under fluoride stress, ComX reached a similar peak protein abundance, but, in contrast to heat stress, its level did not return to baseline. Although its abundance decreased somewhat, ComX remained nearly 10-fold higher than the pre-treated cells at the 30- and 45- minute timepoints (**Figure 3a).**

**Figure 3.**
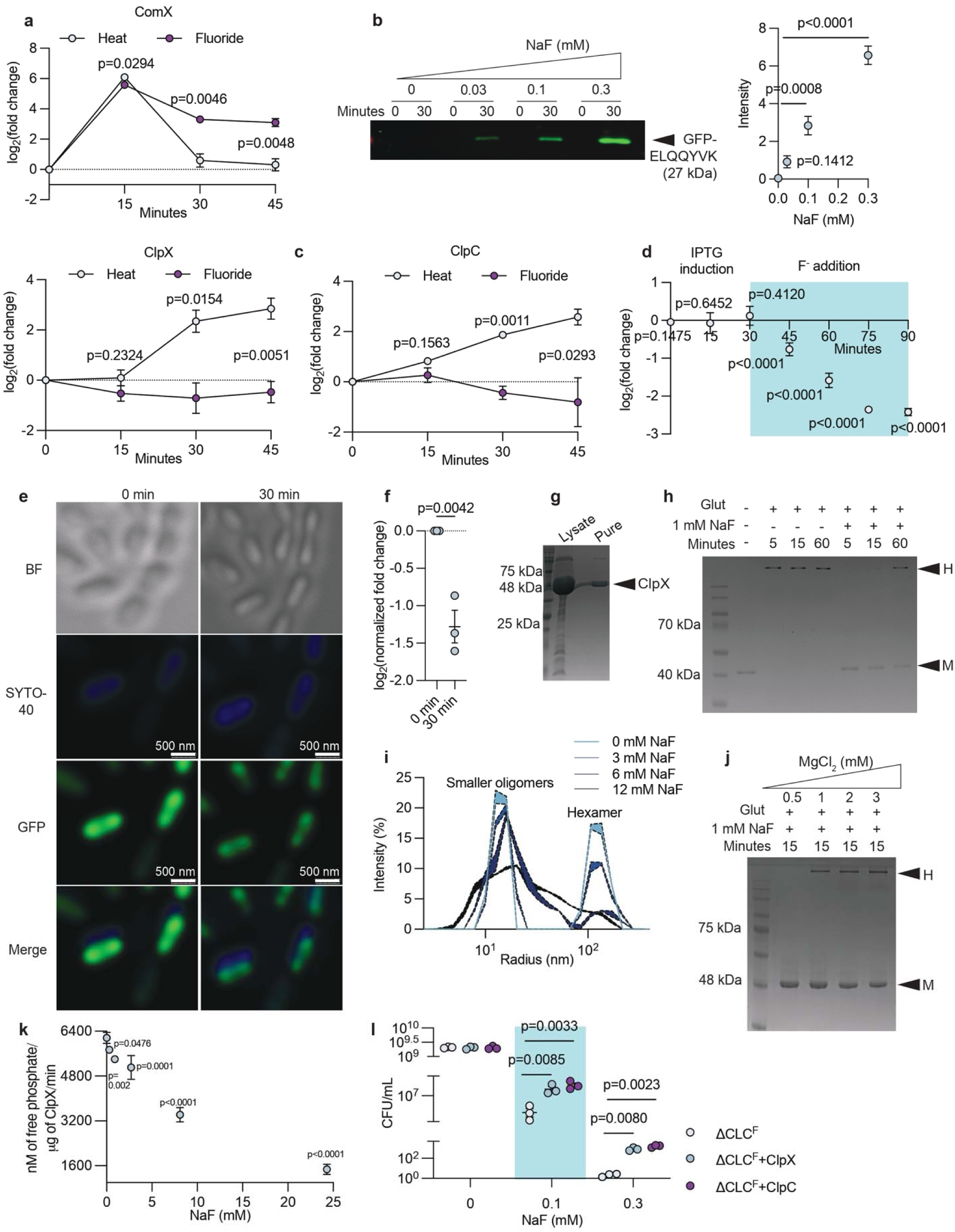
Fluoride disrupts degradation of ComX by Clp protease complex. **a.** Relative ComX abundance in heat- or 0.3 mM NaF-treated cells as a function of time, determined using quantitative mass spectrometry. Data is normalized relative to protein abundance prior to treatment. Datapoints represent mean and SEM of three independent biological replicates (*n*=3). Significance for each timepoint is calculated using two-tailed t-test. **b.** Left, representative immunoblot with detection of GFP in cells treated with increasing NaF for the time indicated. Right, band intensity after 30 min. Datapoints represent mean and standard error for three independent biological replicates. Significance is calculated using one-way analysis of variance (ANOVA) followed by Fisher’s LSD test. The full immunoblot is shown in **Supplementary Figure 8**. **c.** Relative ClpX and ClpC abundance as a function of time, from the same samples as panel **a,** determined using mass spectrometry and quantified as in panel **a**. **d.** Bimolecular fluorescence complementation (BiFC) in cells for ClpP and ClpX tagged with split GFP. Fluoride is added at 30 min (shaded region), with fluorescence monitored as a function of time. At each timepoint, values are normalized against control cells that are not treated with fluoride. Datapoints represent the mean and SEM of nine independent biological replicates (*n*=9). At each point, significance relative to the no-fluoride control is calculated using a two-tailed t-test. **e.** Single cell BiFC using ClpP and ClpX tagged with split GFP as in panel **d,** before and after treatment with 0.3 mM NaF. Cells are stained with SYTO-40 to detect total nucleic acid, and imaged with emission peaks at 509 nm (GFP) and 441 nm (SYTO-40). **f.** Mean BiFC in single cell imaging after fluoride treatment. Each datapoint represents the total GFP fluorescence from three fields of view normalized against the total SYTO-40 blue fluorescence from the corresponding field of view and the 0 min-timepoint (*n*=3). The three datapoints were derived from three different biological replicates. Mean and SEM are shown, and significance is calculated using a two-tailed t-test. **g.** Coomassie-stained SDS-PAGE of samples from the heterologous expression and purification of ClpX (46 kDa). **h.** SDS-PAGE of purified ClpX crosslinked with glutaraldehyde in presence or absence of 1 mM NaF as a function of time. Samples contain 0.5 mM MgCl_2_ and 0.5 mM ATPγS. Bands at the ‘M’ position are consistent with monomers (46 kDa) and those at the ‘H’ position are consistent with hexamers (276 kDa). **i.** Dynamic light scattering (DLS) size distribution of purified ClpX as a function of NaF concentration. Samples contain 0.5 mM MgCl_2_ and 0.5 mM ATPγS. For each trace, the shaded envelope represents the mean ± the standard error determined from three independent experiments (*n*=3). **j.** SDS-PAGE of purified ClpX crosslinked with glutaraldehyde for 15 minutes in the presence of 1 mM NaF and 0.5 mM ATPγS as a function of MgCl_2_ concentration. Bands at monomeric and hexameric positions are labelled. **k.** Free phosphate generated by purified ClpX as a function of NaF, detected using a malachite green colorimetric assay. Reactions were initiated with 1 mM ATP and contain saturating MgCl_2_ (8 mM). Datapoints represent mean and SEM of three independent experiments (*n*=3). Significance relative to the zero-fluoride value is calculated using two-way analysis of variance (ANOVA) followed by Fisher’s LSD test. **l.** CFUs on recovery plates after treatment with 0, 0.1 or 0.3 mM NaF for control cells (ΔCLC^F^ *S. mutans*) or cells overexpressing ClpX or ClpC. Mean and SEM of three independent biological replicates is shown (*n*=3). Significance is calculated using two-way analysis of variance (ANOVA) followed by Fisher’s LSD test (non-significant comparisons are not shown).

ComX degradation is mediated by ClpP protease^50–52^. This dodecameric protease associates as a multimeric complex with a hexameric AAA+ family ATPase, such as ClpX, ClpC, or ClpE. The ATPases recognize, deliver, and unfold client proteins^53^. In particular, in *Streptococci*, ClpX plays a significant role and is an essential partner of ClpP under various stressors^54, 55^. To test the idea that ComX levels remain high because cells experience a general defect in Clp-mediated proteolysis during fluoride stress, we monitored cellular levels of GFP tagged with a C-terminal peptide, ELQQYVK, that specifically targets GFP to ClpX for degradation^56^. Upon fluoride addition, we observed a dose-dependent accumulation of the tagged GFP, indicating that ClpXP function is impaired under fluoride stress (**Figure 3b; loading control Supplementary Figure 8**). This observation provides a potential rationale for the elevated cellular levels of ComX beyond the normal competence window.

Changes in Clp ATPase abundance, assembly, or activity could, in principle, contribute to this diminished protease function under fluoride stress. We sought to test each of these possibilities in turn. We first examined the proteomic abundance of ClpX, ClpC, and ClpE under heat and fluoride stress using quantitative mass spectrometry. Under heat stress, the abundance of ClpC and ClpX increases by a modest factor of ∼2-3 fold. Under fluoride stress, the amount of ClpX and ClpC remains constant, at the same level as control cells (**Figure 3c)**. Thus, the reduction in ClpX mediated cleavage of GFP with increasing fluoride relative to the control cells is not explained by a reduction in ClpX or ClpC levels. ClpE did not exhibit any difference in abundance between heat and fluoride stress, with reduced abundance over time under both conditions (**Supplementary Figure 9**).

We next examined whether assembly of the ClpXP complex is impacted by fluoride. To test assembly of the complex in live cells, we used bimolecular fluorescence complementation, in which ClpP was tagged with the N-terminal portion of GFP, and ClpX was tagged with the C-terminal portion. Without fluoride treatment, we observe fluorescence, consistent with formation of the ClpXP complex. Upon addition of 0.3 mM NaF, we observe a decrease in fluorescence over time, suggesting disassembly of the ClpXP complex (**Figure 3d**). The time-dependent loss of fluorescence was not observed in the absence of fluoride. Live single cell imaging supported the results of the bulk assays, with a significant reduction in GFP fluorescence after 30 minutes with 0.3 mM NaF (**Figure 3e**, **Figure 3f**).

To more directly assess the effects of fluoride on ClpX function, we heterologously expressed and purified *S. mutans* ClpX (**Figure 3g**). We first examined oligomerization of the ClpX hexamer in the presence of fluoride using chemical crosslinking and SDS PAGE (**Figure 3h**). For these experiments, we included the nonhydrolyzable ATP analog ATPγS and Mg^2+^, which have been shown to promote assembly or stability of other bacterial ClpX homologs^57–59^. In the absence of fluoride, ClpX was rapidly crosslinked to the hexameric position of the fully assembled ClpX complex. In contrast, fluoride addition favored a monomeric population, with the hexameric crosslinked species only accumulating after a one-hour incubation. To confirm this observation, we complemented the crosslinking study with a biophysical analysis of complex assembly using dynamic light scattering (DLS, **Figure 3i**), which measures the hydrodynamic radius of particles in solution. As in our crosslinking study, when the fluoride concentration was increased, we saw a reduction in particles consistent with a hexameric species. This was accompanied by a broadening distribution of smaller particles, suggesting an increasingly heterogenous population of incompletely assembled ClpX particles with increasing fluoride.

Because ATP binding and hydrolysis is dependent on Mg^2+^, and ATPγS is associated with oligomerization^58, 60^, we hypothesized that fluoride inhibits ClpX assembly by sequestering Mg^2+^. Indeed, both the crosslinking and DLS analyses showed that the fluoride-related assembly defect could be rescued by the addition of excess Mg^2+^ (**Figure 3j, Supplementary Figure 10**). Moreover, chelation of Mg^2+^ with EDTA exhibited a similar effect on ClpX oligomerization as fluoride addition (**Supplementary Figure 11**), further supporting Mg^2+^ sequestration as the molecular basis for the ClpX assembly defect.

Finally, because fluoride is well known to inhibit the enzymatic activity of ATPases^61–64^, we examined ATPase activity in the presence of saturating levels of Mg^2+^, at which the ClpX hexamer is expected to be fully assembled (**Figure 3k**). These experiments show a fluoride dose-dependent reduction in ATPase activity, although the inhibitory concentrations of fluoride are somewhat higher than those required to promote hexamer disassembly.

Thus, these experiments support a model in which the tight temporal regulation of ComX is undermined by defects in ComX degradation by Clp proteases, mainly ClpXP. By sequestering Mg^2+^, fluoride induces disassembly of the active ClpX hexamer, which contributes to the diminished formation of the ClpXP complex. Fluoride also has a secondary effect of inhibiting ATP hydrolysis by ClpX, even in the presence of saturating Mg^2+^. These results imply that fluoride-induced lysis could potentially be rescued by overexpression of the Clp ATPases. Indeed, plasmid-based overexpression of ClpX or ClpC both exhibited partial rescue of fluoride-induced cell death at 0.1 and 0.3 mM NaF, with >2 orders of magnitude greater survival at the higher fluoride concentration (**Figure 3l**).

### Aromatic amino acids counteract the inhibition of ClpXP-dependent proteolysis by fluoride

Though many genes found in fluoride-responsive operons have well-established roles in the fluoride stress response, there remain some genes whose mechanistic relationship to fluoride stress is unclear. A prominent example is chorismate mutase^16^. A survey of streptococcal species from the Joint Genome Institute’s representative collection of bacterial genomes^65^ reveals that nearly 90% of *Streptococci* possess a second copy of this gene 5’ to the fluoride exporter (**Figure 4a**). Additionally, increased expression of a gene encoding chorismate mutase (*smu_1291*) together with the adjacent CLC^F^-encoding genes has been reported previously in wildtype *S. mutans*^66^, a result that we replicated here (**Supplementary Figure 12**).

**Figure 4.**
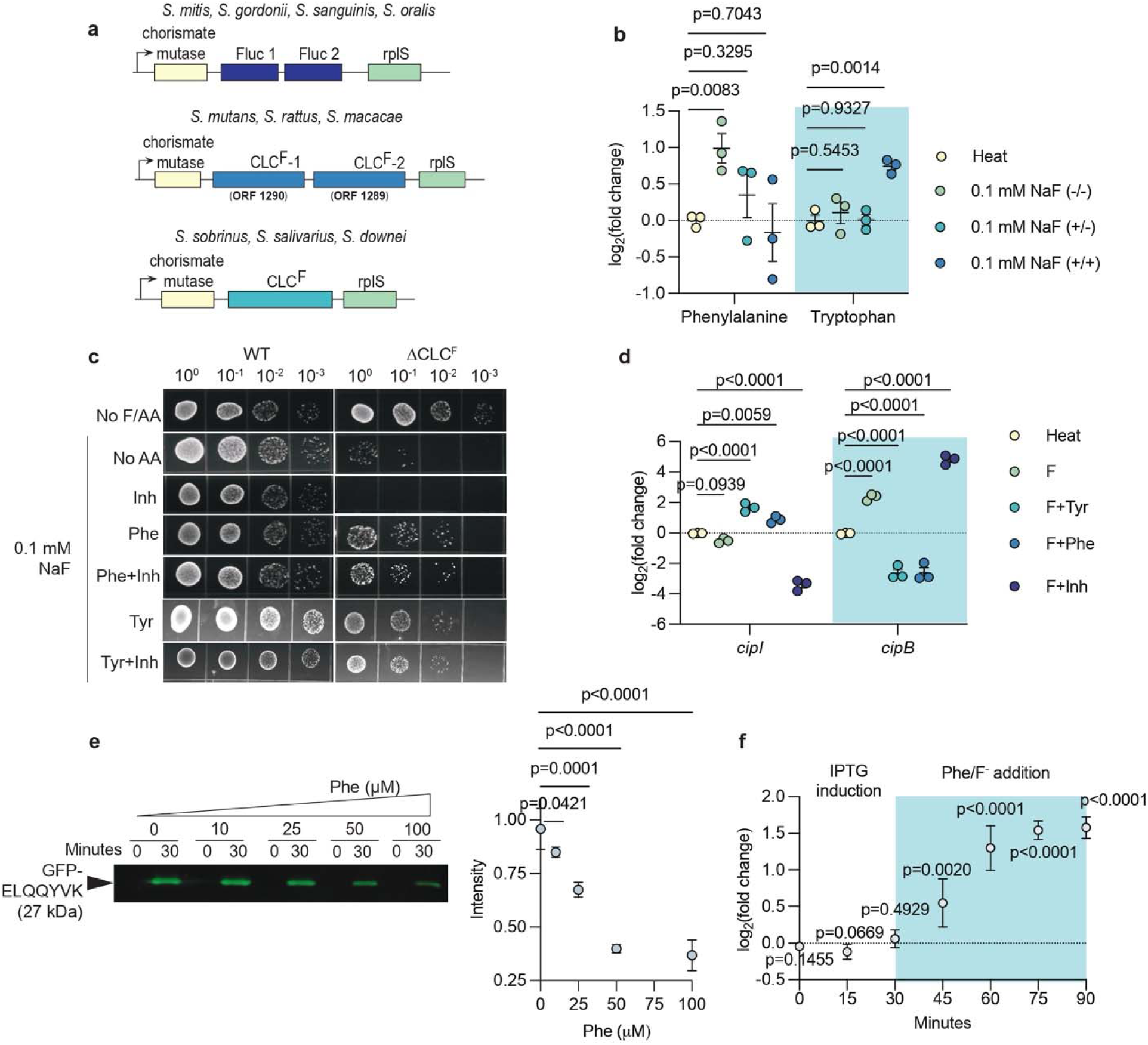
Aromatic amino acids counteract fluoride inhibition of Clp mediated proteolysis. **a.** A gene encoding chorismate mutase is present in fluoride export operons in most *Streptococci*. **b.** Metabolomic analysis of phenylalanine and tryptophan in heat-treated cells and the sorted populations of fluoride-treated cells (**Figure 1d**). Data are normalized with respect to the heat-treated reference sample. **c.** 10-fold serial dilutions of WT and ΔCLC^F^ *S. mutans* with fluoride (0.1 mM), phenylalanine (10 µM), tyrosine (10 µM) or the chorismate mutase inhibitor adamantane-1-acetic acid (Inh, 200 µM) added as indicated. A no-fluoride control is shown in **Supplementary Figure 13**. **d.** Expression of *cipI* and *cipB* in heat- or 0.1 mM NaF-treated cells with added Phe (10 µM), Tyr (10 µM) or adamantane-1-acetic acid (Inh, 200 µM). Expression is normalized relative to the heat-treated reference sample. For **b** and **d**, the datapoints represent mean and SEM of three independent biological replicates (*n*=3). Significance is calculated using two-way analysis of variance (ANOVA) followed by Fisher’s LSD test. **e.** Left, representative immunoblot detection of GFP tagged with the ELQQYVK peptide in lysates of cells treated with 0.3 mM NaF as a function of phenylalanine concentration. Right, band intensities after 30- minute treatment. Data represent the mean and standard error from three independent immunoblots (*n=*3). Intensity is normalized relative to the sample with no added Phe. Significance is calculated using one-way analysis of variance (ANOVA) followed by Fisher’s LSD test. The full immunoblot is shown in **Supplementary Figure 14. f.** Bimolecular fluorescence complementation (BiFC) to detect ClpX and ClpP association as a function of time in cellular lysates treated with 0.3 mM NaF and 10 µM Phe. At each timepoint, data is normalized relative to fluoride-treated cells with no added Phe. Datapoints represent mean and SEM of nine independent biological replicates (*n*=9). At each point, significance relative to the fluoride-treated control with no added Phe is calculated using a two-tailed t-test.

Chorismate mutase is the rate-limiting enzyme of the shikimate pathway, where it catalyzes the conversion of chorismate to prephenate. This step represents the committed step for the biosynthesis of aromatic amino acids phenylalanine and tyrosine, and shunts the metabolic flux away from tryptophan synthesis. To assess whether aromatic amino acids might be correlated with fluoride tolerance, we examined the amino acid pools in the sorted cell populations exposed to 0.1 mM NaF (introduced in Figure 1). We observed significantly higher phenylalanine in the -/- sub-population, which does not exhibit cell wall damage, relative to the +/- or +/+ populations. In contrast, the inviable +/+ sub-population had significantly more tryptophan (**Figure 4b**). Tyrosine was below the limit of detection in all samples. This observation suggests that chorismate mutase-associated flux of amino acid synthesis towards phenylalanine, and away from tryptophan, is associated with limiting cell wall damage and survival under fluoride stress.

Likewise, we also found that addition of exogenous phenylalanine or tyrosine improves the survival of the ΔCLC^F^ *S. mutans* strain at 0.1 mM NaF (**Figure 4c, Supplementary Figure 13**). In contrast, addition of the chorismate mutase inhibitor adamantane-1-acetic acid^67^ aggravated this strain’s susceptibility to 0.1 mM NaF, an effect that was mitigated by additional exogenous phenylalanine. The expression of *cipI* and *cipB* coincided with the results from the cellular resistance assays: Phe or Tyr addition yielded significantly increased expression of the immunity protein-encoding gene *cipI*, and significantly decreased expression of the lysis-activating *cipB* **(Figure 4d)**. The opposite expression pattern was observed upon addition of adamantane-1-acetic acid. These results explicitly link chorismate mutase activity and its downstream products to the competence pathways implicated in fluoride-dependent cell lysis.

We next wondered if phenylalanine can rescue the proteolysis defects that underlie the runaway competence pathways. We revisited the assay monitoring degradation of ClpX-targeted GFP, this time in cell lysates, which allowed us to directly add Phe without depending on cellular uptake mechanisms (**Figure 4e, loading control Supplementary Figure 14**). This experiment showed restoration of ClpXP-mediated proteolysis of the GFP test substrate by phenylalanine in a dose-dependent fashion. We also assessed ClpXP assembly using bimolecular fluorescence complementation in live cells, which showed that 10 μM Phe partially restored formation of the ClpXP complex (**Figure 4f**). This was not due to any impact on ClpX oligomerization, which was not altered by phenylalanine addition *in vitro* (**Supplementary Figure 15**). Together, these results show that the increased synthesis of aromatic amino acids counteract the prolonged autolytic signaling in the fluoride stressed cells, with a concomitant partial restoration of assembly and activity of ClpXP protease.

## Discussion

In the present work, we uncover the molecular mechanism of fluoride-mediated bacterial killing, nearly 50 years after Marquis first described fluoride as causing “massive lysis” of several different species^25^. Whereas mechanisms of fluoride growth inhibition are well understood^20, 29^, our findings represent the first molecular description of fluoride-induced cell death.

Our work establishes that, in *S. mutans,* the widespread lysis is due to a runaway stress response caused by fluoride’s inhibition of the proteolytic machinery (**Figure 5**). While limited autolysis of a sub-population is an adaptive mechanism that promotes gene transfer to sister cells under starvation or stress^36–38, 68^, this mechanism is normally under tight temporal control^49, 50^. By sequestering Mg^2+^, fluoride causes disassembly of the Clp ATPases, undermining the cell’s ability to degrade the competence-associated alternative sigma factor ComX, and prolonging the lytic response. Our findings are bolstered by a previous observation that ClpP is important for survival of *S. mutans* under fluoride stress^69^.

**Figure 5.**
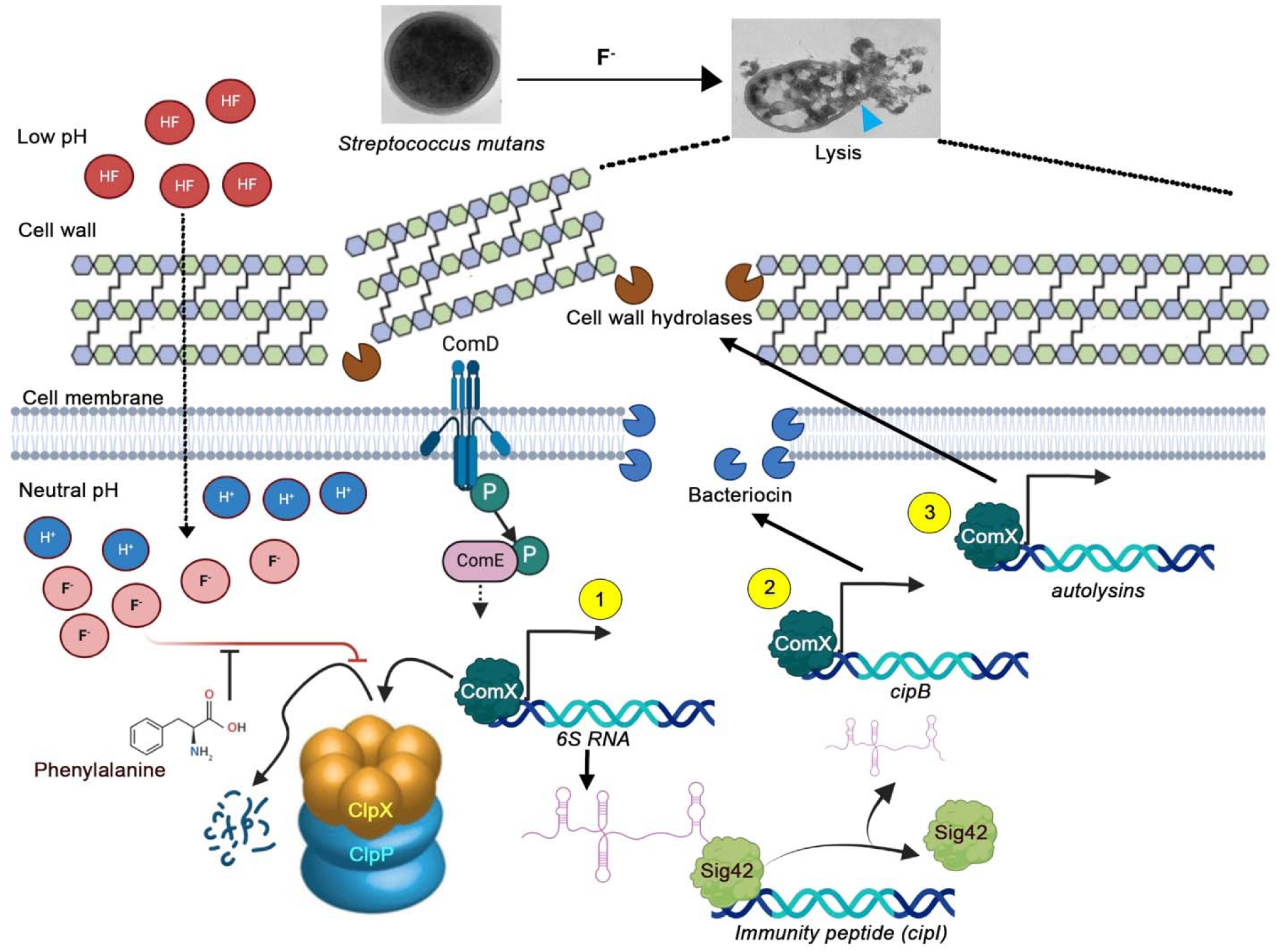
Model for fluoride-dependent cell lysis in *Streptococcus mutans.* Fluoride accumulation inhibits the oligomerization, assembly and ATPase activity of ClpX required to power the ClpP-proteolytic complex. This prolongs the expression window of the competence-associated alternative sigma factor ComX, a client of ClpXP. ComX induces the expression of (1) 6S RNA, which dissociates the primary sigma factor Sig42 from the promoter of the immunity peptide (*cipI*); (2) bacteriocins, like *cipB*; and (3) autolysins. Fluoride-induced expression of chorismate mutase and resultant phenylalanine synthesis partially restores ClpXP activity. TEM micrographs showing cell lysis are adapted from Banerjee et al. ^4^.

Thus, fluoride’s bactericidal effect shares a root cause with fluoride’s bacteriostatic effects: the disruption of metal homeostasis. Bacteriostatic inhibition mechanisms primarily target the activity of metalloenzymes, either because fluoride sequesters active site metal ions, or because it forms fluoride-metal complexes that act as transition state inhibitors^20^. In the case of the Clp ATPases, fluoride interferes not only with enzymatic activity, but also with oligomerization, which we show is Mg-ATP-dependent in *S. mutans.* The cell wall damage caused by sustained ComX signaling and the expression of cell wall hydrolases is not as easily reversible as metabolic inhibition.

Fluoride has been observed to cause bacterial cell lysis in other diverse species as well, in this study (**Supplementary Figure 2**) and others^4, 25–27^. The mechanism of cell death described here requires both a stress-induced lytic response, and a fluoride-sensitive turnoff mechanism. We expect this mechanism to be general among *Streptococci*, in which ComX-dependent competence and its degradation by Clp proteases are conserved^49–52, 70^. Moreover, autolytic stress responses are widespread among bacteria^71^. Although not all bacteria rely on Clp proteases to terminate signaling, those that do are likely to be sensitive to perturbations to metal homeostasis, since a requirement for a divalent metal is a common theme in Clp protease assembly, stability, and activity^53, 57–59, 72^. Thus, it is likely that analogous mechanisms underlie fluoride-induced cell death in other bacterial genera as well.

Finally, this work also addresses another persistent question about the bacterial fluoride response: the role of aromatic amino acids in counteracting fluoride toxicity. The association between chorismate mutase and fluoride-responsive operons was originally noted when these operons were first discovered^16^, and increased chorismate mutase expression and cellular aromatic amino acid pools have been observed previously in fluoride-stressed microbes^9, 21^. We show that phenylalanine promotes *in vivo* ClpXP assembly, and partially restores Clp-dependent proteolysis during fluoride stress. A terminal phenylalanine is a common feature of the various cellular effectors and peptides that influence Clp protease assembly and activity in different bacterial systems^73, 74^, and we speculate that free phenylalanine might access this binding pocket as well. It has also been suggested that aromatic amino acids mitigate oxidative stress, which often accompanies fluoride stress^9^. However, the present experiments suggest that, at least in the *Streptococci*, the products of chorismate mutase play a more direct role in the cellular response to fluoride stress.

In summary, we show that fluoride causes the misregulation of the tightly controlled competence signaling pathway, promoting cell lysis in *S. mutans,* the etiological agent of dental caries. This phenomenon implies that, at least for some bacteria, antimicrobials could be developed that potentiate fluoride’s bactericidal effects. This study reveals several such targets, including fluoride efflux pumps, chorismate mutase, and the Clp protease complex.

## Methods

### Bacterial strains, media and growth conditions

Bacterial strains used in this study are listed in **Supplementary Table 1**. *Escherichia coli* was cultured aerobically at 37°C in Luria Bertani media (LB). Typically, *L. salivarius* and *Streptococci* were cultured in brain heart infusion media (BHI) with 1% glucose (w/v) at 5% CO_2_ and 37°C. Cultures of ΔCLC^F^ *S. mutans* also included 0.3 mg/mL spectinomycin and 10 µg/mL erythromycin. For ClpX and ClpC overexpression strains, synthetic genes (Integrated DNA Technologies, USA) were inserted into pIB184-Km under a constitutive P23 promoter^75^ and used to transform ΔCLC^F^ *S. mutans*, which was cultured with 0.5 mg/mL kanamycin.

### Calibration of fluoride, heat, and chloramphenicol treatment

Cultures were grown until an OD_600_ of 0.5 at 37°C, and re-inoculated in fresh media containing 60 mM NaF to a final OD_600_ of 0.1. Cultures were exposed to heat, fluoride, or chloramphenicol stress and then re-plated on BHI recovery plates containing 1% (w/v) glucose. CFUs were counted after 48h incubation. The chloramphenicol and heat exposures were increased incrementally until conditions were identified that yielded comparable CFUs after recovery as treatment with 0.3 mM NaF for 4 hours. These conditions are 52°C for 45 min for the heat treatment, or 10 µg/mL chloramphenicol for 4 hours.

### Transmission electron microscopy (TEM)

After treatment, cultures were pelleted and fixed with glutaraldehyde (2% in 0.1 M Sorensen’s buffer, pH 7.4) for 16h at 4°C. After washing, samples were incubated in 1% osmium tetroxide for 2h at 25°C. Cells were immobilized in 1% agarose and ∼0.5 mm slices were cut and dehydrated in ethanol (four 15-minute washes, 25% −100%) and washed with propylene oxide. A resin composed of Poly/Bed812 (PolySciences Inc., USA), dodecenylsuccinic anhydride (DDSA), and nadic methyl annhydride (NMA) in a 2:1:1 ratio in propylene oxide, with 2% Tris(dimethylaminomethyl)phenol (DMP-30) as an accelerator, was used to infiltrate the dehydrated agarose slices in three concentration steps (25%, 50%, and 75%, 16 hours per step) prior to a final incubation in electron microscopy molds (60°C, 48h under vacuum). After polymerization, samples were sectioned (50 nm) with an ultramicrotome (Leica UC7), loaded onto copper grids, stained with 7% (w/v) uranyl acetate, washed in deionized water, and post-stained in Reynold’s lead citrate. Samples were imaged using a JEOL 1400-plus transmission electron microscope equipped with an XR401 AMT sCMOS camera.

### Quantitative PCR

Total RNA was isolated from *S. mutans* cell lysate using TRIzol Max Bacterial RNA isolation kit (Thermofisher Scientific, USA) using the manufacturer’s protocol and treated with DNAseI to remove genomic DNA contamination. Reverse transcription was performed using Maxima First Strand cDNA synthesis kit (Thermofisher Scientific, USA) following the manufacturer’s protocol. For qPCR, gene-specific primers (**Supplementary Table 2**) and 10 ng of cDNA were used. Amplification was monitored on the qPCR Step-One Plus detection system (Applied Biosystems, USA) using Power SYBR Green PCR Master Mix (Thermo Fisher Scientific, USA). Expression variance of the target genes was calculated based on the 2^-ΔΔC^ formula using expression of endogenous control *gyrA*.

### Fluorescence-activated cell sorting (FACS)

Cells were stained with AnnexinV-FITC and propidium iodide (PI) (Invitrogen, USA) following the manufacturer’s protocol. The cells were sorted using a Bigfoot Spectral Cell Sorter (Thermo Fisher Scientific, USA) with λ_ex_ 561 and 488 nm, and emission band-passes 625±15 and 507±19 nm for PI and FITC, respectively. Sorting was performed at a low flow rate of 20 psi, voltage settings of 525 V (forward scatter), 630 V (side scatter), 650 V (PI) and 610 V (FITC) and in a 2D trigger mode using 0.58% (side scatter) and 0.25% (forward scatter) as thresholds. ∼100,000 cells were collected from each fraction for downstream analyses. Data were visualized and analyzed using FlowJo (BD Bioscience, USA).

### Promoter pull-down

Streptavidin Magnesphere Paramagnetic particles (Promega, USA) beads were washed and resuspended in 10 mM Tris-HCl (pH 7.5), 1 mM EDTA and 2 M NaCl. Promoter sequences (**Supplementary Table 3**) were PCR amplified using 5’-TEG-biotin labelled primers (Integrated DNA Technologies, USA) (**Supplementary Table 2**), and the amplicons were purified and concentrated using the Monarch Spin PCR & DNA Cleanup Kit (New England Biolabs, USA) and incubated with the magnetic beads (1 mg/200 µL amplicon) for 4h. The supernatant was discarded, and beads were rinsed with wash buffer, TE buffer (0.5 M Tris-HCl pH 8, 1 mM EDTA), BS/THES buffer (22 mM Tris-HCl pH 7.5, 4.4 mM EDTA, 8.9% sucrose, 62 mM NaCl, protease inhibitor cocktail, 2 mM HEPES pH 7.5, 1 mM CaCl_2_, 10 mM KCl, 2.4% glycerol) and BS/THES buffer containing salmon sperm DNA. Cell lysates, pre-treated with streptavidin-conjugated agarose beads to remove non-specific binders, were incubated with the magnetic beads at 4°C for 16 h. The beads were immobilized and rinsed well with fresh BS/THES buffer prior to elution in high salt buffer (25 mM Tris-HCl pH 7.5, 0.5 M NaCl) and analysis by SDS-PAGE. For protein identification, bands from the gel were excised, crushed and extracted with 50 mM ammonium bicarbonate pH 8 and 8 M urea for 16h. Supernatants were precipitated using chilled acetone at −80°C for 3h and centrifuged at 15,000 × g to pellet the protein. The protein pellet was washed in acetonitrile, dried, and re-suspended in 50 mM ammonium bicarbonate pH 8 with 2 M urea, and digested overnight at 37°C with Trypsin Gold (Promega, USA) in a final protease:protein ratio of 1:50. The reaction was terminated using 1% formic acid. Peptides were desalted and reconstituted in 2% acetonitrile and 0.1% formic acid. High-energy collisional dissociation MS/MS spectra were acquired on an Orbitrap Fusion Lumos mass spectrometer (Thermofisher Scientific, USA) and peptide and protein identification was performed with Sequest on Proteome Discoverer 2.2 (Thermofisher Scientific, USA) against a composite of forward/reverse sequences of NCBI-*Streptococcus mutans* UA159 (NC_004350.2) and common lab contaminants. A 1% false discovery rate (FDR) was applied to filter peptide and protein identification. Protein abundance was calculated from the sum of peptide ion abundances based on ion intensity. For RNA identification, the RNA fraction was separated using agarose gel electrophoresis and reverse transcribed using oligodT primers with the Maxima First Strand cDNA synthesis kit (Thermofisher Scientific, USA) according to the manufacturer’s protocol, followed by sequencing using oligodA primers.

### Biolayer interferometry (BLI)

Biolayer interferometry experiments were conducted using an OctetRed instrument (Sartorius, USA). The biotinylated amplicons of *cipI* and *cipB* promoters were immobilized on streptavidin-conjugated sensor tips. For association, the tips were incubated with the eluted fractions from promoter pull-down assays. For dissociation, the tips were incubated in elution buffer, 25 mM Tris-HCl pH 7.5, 0.5 M NaCl. BLI data were visualized using the OctetRed Data Analysis software (Sartorius, USA).

### Electrophoretic mobility shift assay (EMSA)

6S RNA was prepared by *in vitro* transcription using HiScribe T7 High Yield RNA Synthesis Kit (New England Biolabs, USA) and a gBlock gene fragment (Integrated DNA Technologies, USA) with a T7 promoter 5’ to the 6S sequence as the template. 3.5 mM fluorescein-12-UTP (Sigma Aldrich, USA) was added together with 6.5 mM UTP to incorporate the label at approximately every 20 to 25^th^ nucleotide of the transcript. After spin column cleanup of the product, RNA was denatured at 90°C for 3 min in 10 mM sodium phosphate and refolded at 50°C for 20 min with 10 mM MgCl_2_ and 10 mM NaCl. The *cipI* promoter was amplified using primers with Cy5 labels at their 5’-ends (Integrated DNA Technologies, USA). Cellular lysates were suspended in 10 mM sodium phosphate buffer pH 8, 1 mM EDTA, 80 mM MgCl_2_ and 2 µg poly (dI-dC). Lysates were pre-incubated with 10 ng of Cy5-labelled *cipI* promoter for 20 minutes on ice prior to addition of a 10-fold molar excess of fluorescein-labelled 6S RNA. DNA-protein-RNA interactions were visualized on a native 5% polyacrylamide gel using Cy5 and fluorescein emission.

### Proteomics

100 µg of total protein, quantified by Bradford assay, was precipitated using acetone, washed with acetonitrile and dried. Dried protein samples were denatured in 50 mM ammonium bicarbonate pH 8 with 8 M urea, reduced with 5 mM dithiothreitol and alkylated with 10 mM iodoacetamide. Proteins were trypsin digested (Promega, sequencing grade) at a ratio of 1:100 enzyme: protein at 37°C overnight. The reaction was quenched by adding acetic acid pH 3. Peptides were desalted (Pierce C18, Thermofisher Scientific, USA) and reconstituted in sample loading solution (2% acetonitrile with 0.1% formic acid) prior to liquid chromatography-mass spectrometry (LC-MS) analysis. Samples were separated on a C18 column (Acclaim PepMap RSLC) at 300 nL/min over a 120 min gradient of solvent A (2% acetonitrile with 0.1% formic acid) and solvent B (80% acetonitrile with 0.1% formic acid) by nano-UPLC (Ultimate 3000, Thermofisher Scientific, USA). Peptides separated by LC were introduced into an Orbitrap Fusion Lumos MS (Thermofisher Scientific, USA) via positive mode nano-electrospray (ESI). MS analysis on Orbitrap was performed over 375-1600 m/z at 120K resolution. Tandem MS/MS was performed for precursor ions over 5×10^4^ ion abundance with charge state of 2-7. Peptides and proteins were identified from LC-MSMS data as described in the *Promoter pulldown* section. Specific tryptic peptides with a mass tolerance of two mis-cleavages within 10 ppm, fixed modification of cysteine carbamidomethylation, and dynamic modifications like methionine oxidation, asparagine deamidation and protein N-terminal acetylation were allowed in the search. Normalized protein abundances were compared across samples for label-free quantification of differential protein expression.

### Immunoblotting

A codon optimized synthetic gene encoding GFP-ELQQYVK (Azenta, USA) was inserted into a pIB184Km vector under a constitutive P23 promoter and transformed into ΔCLC^F^ *S. mutans.* After cell culture, total protein in the cell lysate was estimated by Bradford assay. Immunoblotting was performed with 15 µg of protein per sample using anti-rabbit GFP primary antibody (Torrey Pines Biolabs, USA) and goat anti-rabbit secondary antibody tagged to IRDye 800CW (LICORbio, USA). Blots were imaged using a LICOR fluorescence scanner. Band intensities from independent blots were analyzed using ImageJ^76^. Protein loading was confirmed by Coomassie-stained SDS-PAGE.

### Bimolecular fluorescence complementation (BiFC)

Split GFP^77^ coding sequences were codon optimized for *S .mutans.* gBlock gene fragments with an N-terminal fragment of GFP fused to ClpP and a C-terminal fragment of GFP fused to ClpX (Integrated DNA Technologies, USA) were inserted into a pDL278 vector under an IPTG-inducible promoter, and a pIB184-Km vectors under a constitutive promoter, respectively. The plasmids were transformed in Δ1289 *S. mutans*, which is phenotypically indistinguishable from ΔCLC^F^ ^4^, and cultured in BHI supplemented with 1% glucose, 0.5 mg/mL kanamycin, 0.3 mg/mL spectinomycin and 10 µg/mL erythromycin.

For bulk experiments, cultures were inoculated in black-bottom 96 well plates and incubated at 37°C without shaking. Expression of the ClpX/C-GFP chimera was induced with 0.5 mM IPTG for 30 min prior to addition of 0.3 mM NaF. GFP fluorescence was measured at 15- minute intervals (λ_ex_=485; λ_em_= 535) using an Infinite F200 Pro plate reader (Tecan, USA). For single cell imaging, cells were stained with 10 µM SYTO-40 for 20 min, washed with phosphate buffered saline pH 7.4, and resuspended in fresh media containing 0.5 mM IPTG. After 30 minutes induction, 0.3 mM NaF was added. Immediately, 5 µL samples were mounted on thin slices of 0.8% low melting point agarose on microscopy slides. Cells were imaged at 63x magnification on a SP8 Laser Scanning Confocal Microscope (Leica Microsystems, Germany) under oil immersion at 37°C. SYTO-40 and GFP were imaged at 405 and 488 nm respectively. Z-stacked images acquired at 0 and 30 min were maximum projected using LAS-X software (Leica Microsystems, Germany) and deconvoluted using Huygens Professional image processing (Scientific Volume Imaging, Netherlands). Total fluorescence intensity from the blue and green channels were separately estimated as integrated density using ImageJ.

### Purification, protein crosslinking, and ATPase activity of ClpX

The coding sequence of ClpX with a C-terminal hexahistidine tag was cloned into a pASK vector^14^ and transformed into *Escherichia coli* (BL21-DE3). Protein expression was induced at OD_600_ 0.5 with 0.2 mg/mL anhydrotetracycline for 16h at 25°C. ClpX was purified from clarified cell lysates using cobalt affinity resin (1mL/L culture, Takara Bio), washed with 100 mM NaCl, 20 mM imidazole, 20 mM Tris-HCl pH 8.0, and eluted with 400 mM imidazole. A final size exclusion purification step (Superdex 200) was performed in 100 mM NaCl, and 20 mM HEPES pH 7.5. 10% glycerol was added immediately after the size exclusion chromatography, along with MgCl_2_, ATPγS or EDTA as indicated. For crosslinking analysis, purified ClpX (0.2 mg/mL) was incubated with 0.125% glutaraldehyde (∼500-fold molar excess) prior to quenching with 0.15 M Tris-HCl pH 7.5 and analysis by SDS-PAGE. ATPase activity of purified ClpX was measured using the ATPase/GTPase Activity Assay Kit (Sigma-Aldrich, USA) following the manufacturer’s protocol.

### Dynamic light scattering

1 mg/mL purified ClpX was diluted in buffer containing 0.5 mM MgCl_2_ and 0.5 mM ATPγS and additional NaF or MgCl_2_ as indicated, and evaluated using a DynaPro Nanostar dynamic light scattering (DLS) instrument (Wyatt Technology Corp., USA). Ten 30 s acquisitions were averaged for each replicate. DLS spectra were visualized and analyzed using DYNAMICS software (Wyatt Technology Corp., USA).

### Central carbon metabolomics

Approximately 100,000 cells from the sorted populations were sonicated and extracted in 8:2 methanol: water containing 1 µM tricarboxylic acid (TCA) as an internal standard. Cell debris were pelleted, and supernatant was transferred to a liquid chromatography-mass spectrometry (LC-MS) autosampler insert, dried under nitrogen, and reconstituted in 8:2 ratio of water:methanol for LC-MS analysis using an Agilent Infinity Lab II UPLC coupled with a 6545 Q-TOF MS in negative ion mode^78^. For quantitative analysis, data were processed using MassHunter Quantitative analysis version B.07.00 (Agilent Technologies, USA). Metabolites were normalized to the nearest isotope labeled internal standard and quantified using a linear calibration curve from replicate injections of Cambridge Isotopes MSKTCA1-US unlabeled TCA calibration standards. Fold change was calculated in log_2_-scale relative to the heat stress.

### Statistics and software

GraphPad Prism10 was used to perform statistical analyses. Data represent mean ± SEM using one- or two-way analysis of variance (ANOVA) with Fisher’s LSD test or two-tailed unpaired t-test, as described in the figure legends.

## Acknowledgements

This work was supported by National Institutes of Health grants 2R35 GM128768 (NIH/NIGMS) to R.B.S. and RM1 DE034220 (NIH/NIDCR) to R.B.S. and L.M.A.T. Orbitrap Fusion Lumos mass spectrometry in the University of Michigan Proteomics Core facility was supported by the Office of the Director, National Institutes of Health under Award Number S10OD021619. Metabolomics measurements were performed by the University of Michigan Metabolomics Core (University of Michigan Medical School, Biomedical Research Core Facilities).

## Author Contributions

Aditya Banerjee: conceptualization, investigation, writing – original draft, visualization; Sophie Pickelner: investigation; Faith Smith: investigation; Livia M. A. Tenuta: conceptualization, writing – review and editing, supervision, funding acquisition, project administration; Randy B. Stockbridge: conceptualization, data visualization, writing – review and editing, supervision, funding acquisition, project administration.

## Supplementary Figures

**Supplementary Figure 1.**
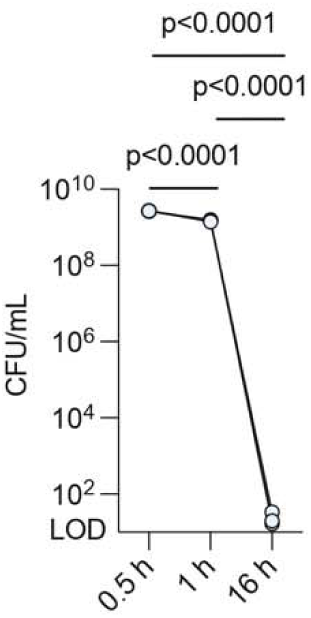
Recovered WT *S. mutans* CFUs on fluoride-free recovery plates after treatment with 60 mM NaF for 0.5, 1 or 16h. Datapoints represent three independent biological replicates (*n*=3). Significance is calculated using one-way analysis of variance (ANOVA) followed by Fisher’s LSD test. LOD stands for limit of detection.

**Supplementary Figure 2.**
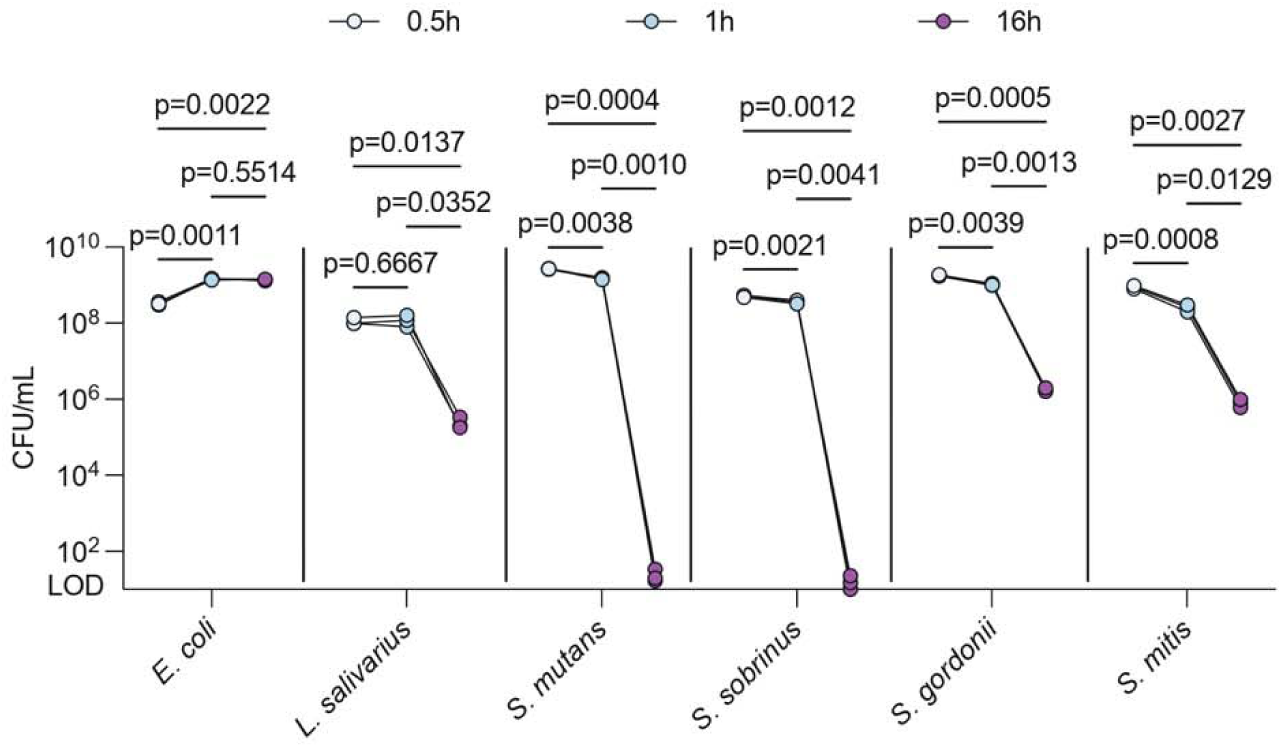
Recovered CFUs on fluoride-free recovery plates after treating WT strains of *E. coli*, *L. salivarius*, *S. mutans*, *S. sobrinus*, *S. gordonii* and *S. mitis* with 60 mM NaF for 0.5, 1 and 16h. Datapoints represent three independent biological replicates (*n*=3). Significance is calculated using two-way analysis of variance (ANOVA) followed by Fisher’s LSD test. LOD stands for limit of detection.

**Supplementary Figure 3.**
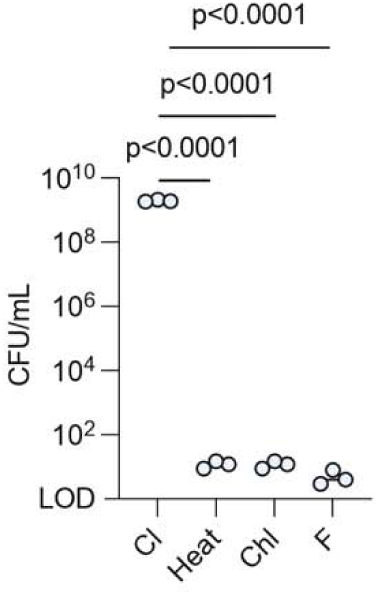
Recovered ΔCLC^F^ *S. mutans* CFUs on recovery plates after treatment with 0.3 mM NaCl or NaF for 4h, 10 µg/mL chloramphenicol (Chl) for 4h or 52° heat for 45 min. Datapoints represent mean and SEM of three independent biological replicates (*n*=3). Significance is calculated using one-way analysis of variance (ANOVA) followed by Fisher’s LSD test. LOD stands for limit of detection.

**Supplementary Figure 4.**
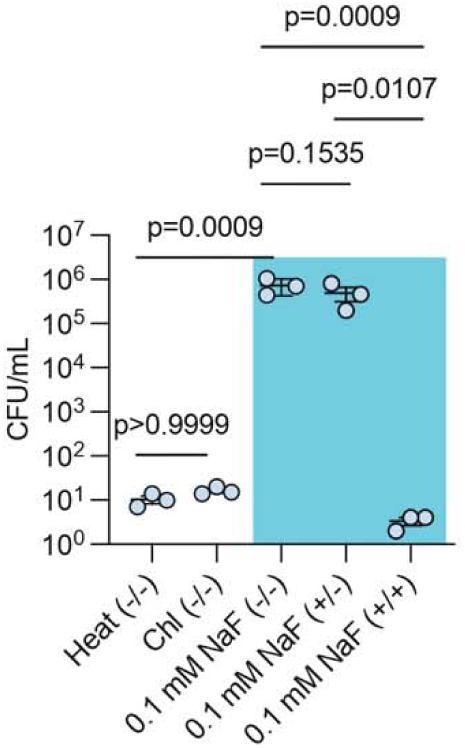
Recovered ΔCLC^F^ *S. mutans* CFUs on BHI plates from the sorted fractions after treatment with heat, chloramphenicol (Chl), or NaF. For heat- and chloramphenicol-treated samples, the -/- fraction is shown. For 0.1 mM NaF, the -/-, +/- and +/+ fractions are shown. Datapoints represent mean and SEM of three independent biological replicates (*n*=3). Significance is calculated using one-way analysis of variance (ANOVA) followed by Fisher’s LSD test.

**Supplementary Figure 5.**
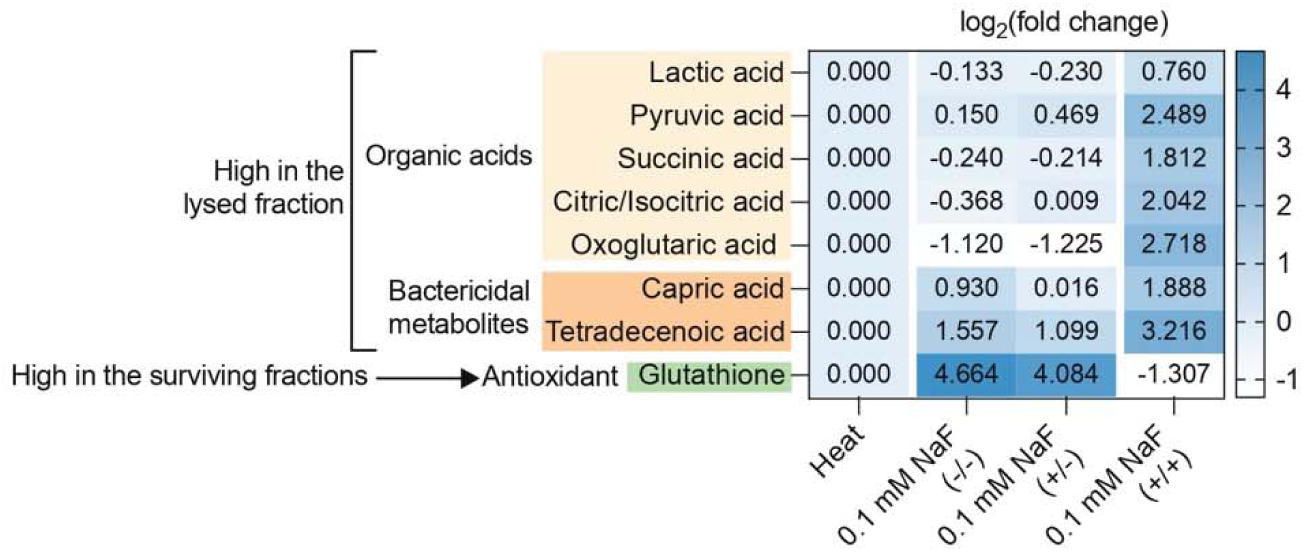
Metabolomics analysis of heat-treated ΔCLC^F^ *S. mutans,* or the sorted fractions of the ΔCLC^F^ *S. mutans* treated with 0.1 mM NaF. Datapoints represent the mean from three independent biological replicates (*n*=3), and fold change has been calculated relative to the heat-treated cells. Significance is calculated using two-way analysis of variance (ANOVA) followed by Fisher’s LSD test.

**Supplementary Figure 6.**
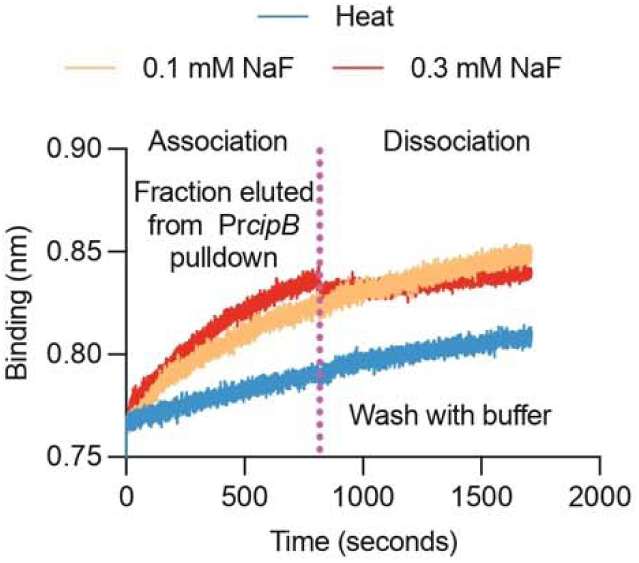
Biolayer interferometry trace showing the association and dissociation phases for biotinylated Pr-*cipB* after incubation with the Pr*cipB* pull-down fractions from ΔCLC^F^ *S. mutans* lysates treated with heat or NaF as indicated.

**Supplementary Figure 7.**
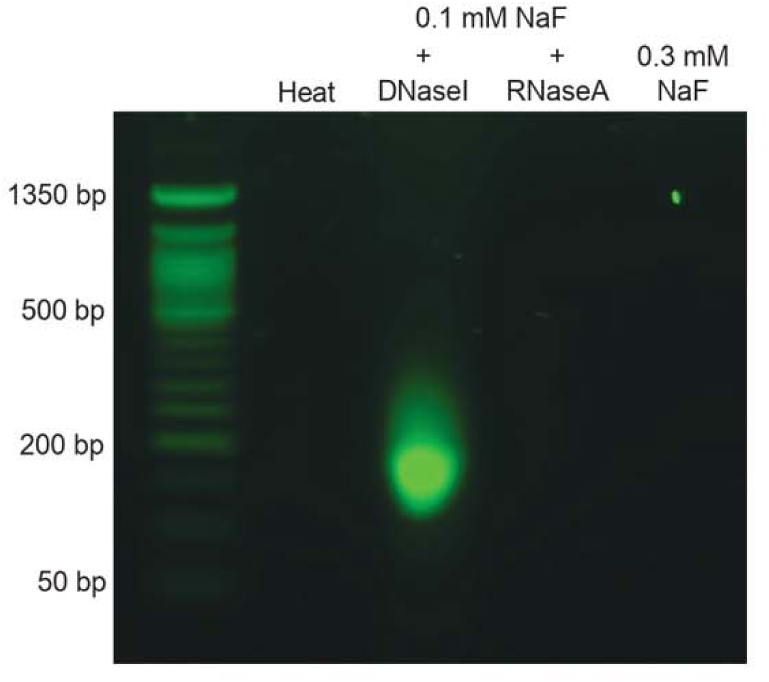
Agarose gel electrophoresis of the Pr-*cipI* pull-down fraction of heat- or NaF-treated ΔCLC^F^ *S. mutans* cell lysates. The fraction eluted from the 0.1 mM NaF-treated cell lysate was further treated with 1 unit of DNaseI or RNaseA prior to electrophoresis.

**Supplementary Figure 8.**
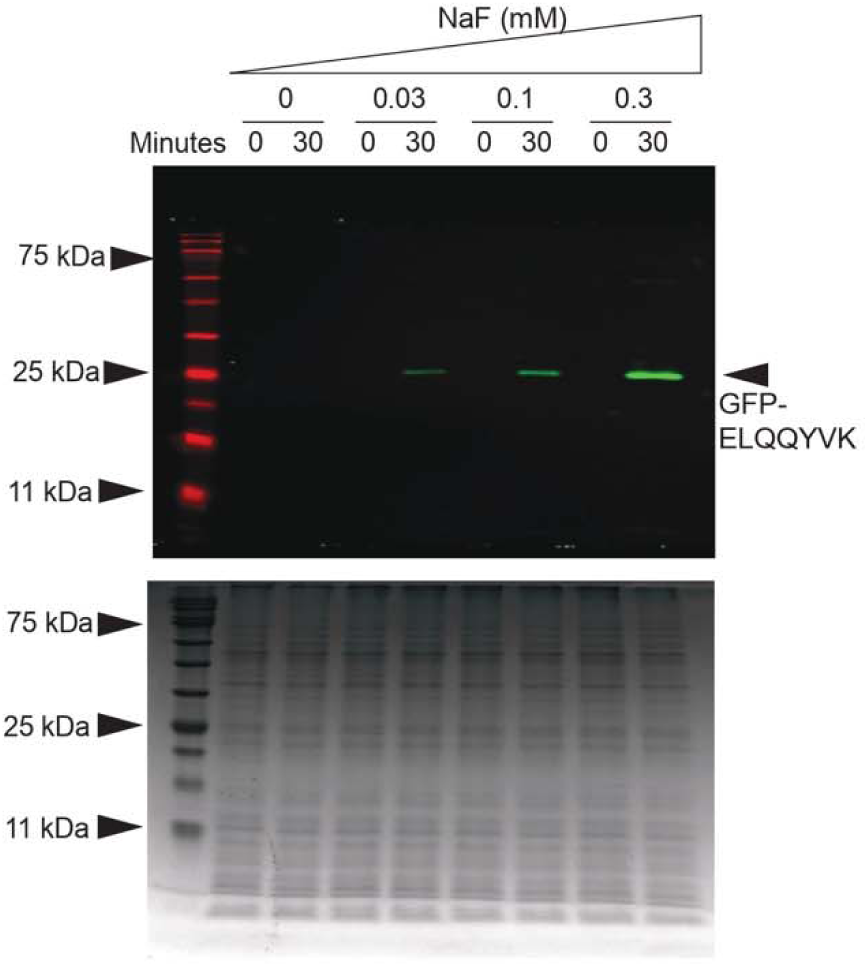
Full immunoblot from Figure 3b, and the Coomassie-stained SDS-PAGE as a loading control.

**Supplementary Figure 9.**
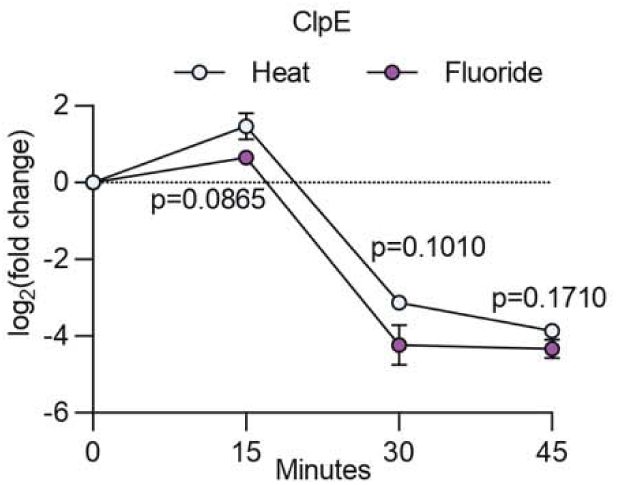
Relative ClpE abundance in ΔCLC^F^ *S. mutans* treated with heat or 0.3 mM NaF as a function of time, determined using quantitative mass spectrometry. Datapoints represent mean and SEM of three independent biological replicates (*n*=3). Significance for each timepoint is calculated using a two-tailed t-test.

**Supplementary Figure 10.**
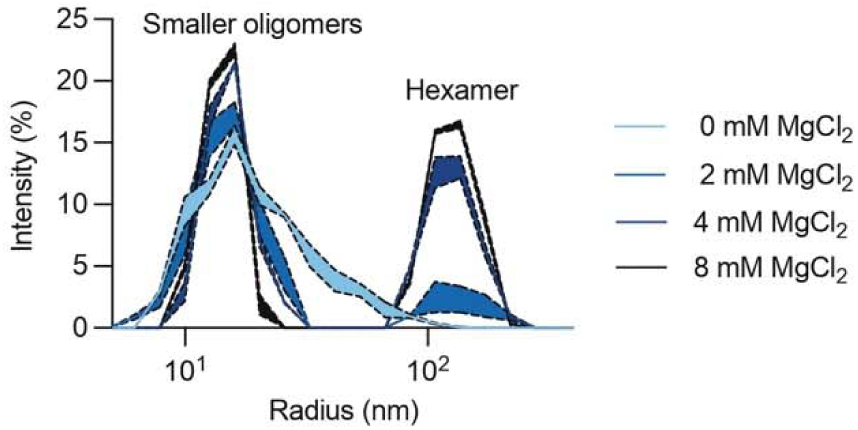
Dynamic light scattering (DLS) of purified ClpX incubated with 3 mM NaF and increasing MgCl_2_. For each trace, the shaded envelope represents the mean ± the standard error determined from three independent experiments (*n*=3).

**Supplementary Figure 11.**
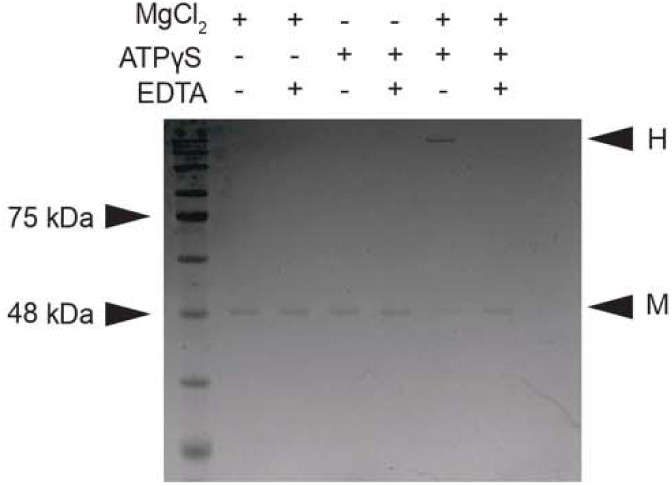
SDS-PAGE of glutaraldehyde-crosslinked ClpX with 1 mM NaF, 0.5 mM MgCl_2_, and 0.5 mM ATPγS or 0.5 mM EDTA, as indicated. Bands labeled ‘M’ are consistent with monomers (46 kDa) and those labelled ‘H’ are consistent with hexamers (276 kDa).

**Supplementary Figure 12.**
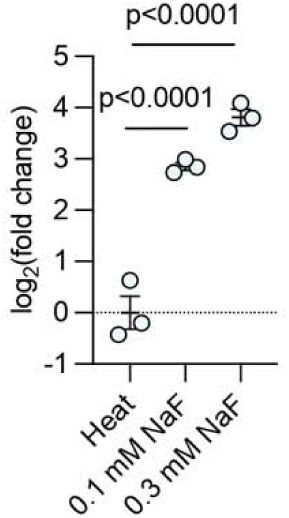
Expression of *smu_1291* (chorismate mutase) in WT *S. mutans* cells after treatment with heat or NaF as indicated. Datapoints represent mean and SEM of three independent biological replicates (*n*=3). Significance is calculated using two-way analysis of variance (ANOVA) followed by Fisher’s LSD test. Gene expression is normalized relative to the heat-treated reference sample.

**Supplementary Figure 13.**
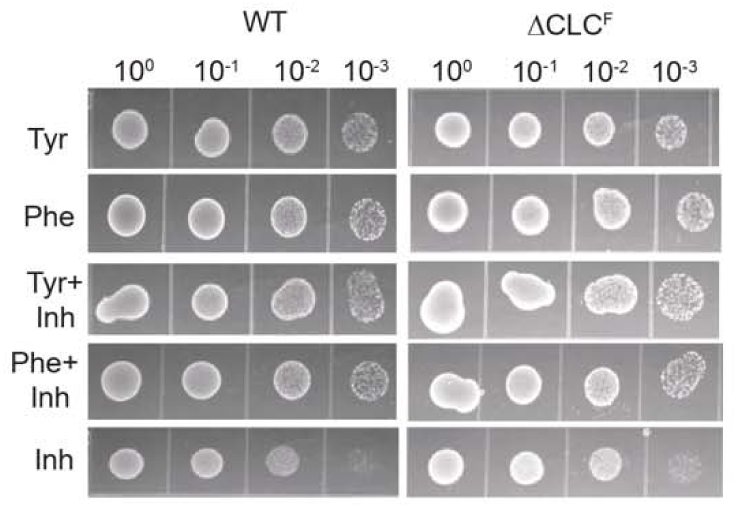
10-fold serial dilutions of WT and ΔCLC^F^ *S. mutans* on fluoride-free BHI plates supplemented with combinations of 10 µM Phe or Tyr and/or 200 µM adamantane-1- acetic acid (Inh) as indicated.

**Supplementary Figure 14.**
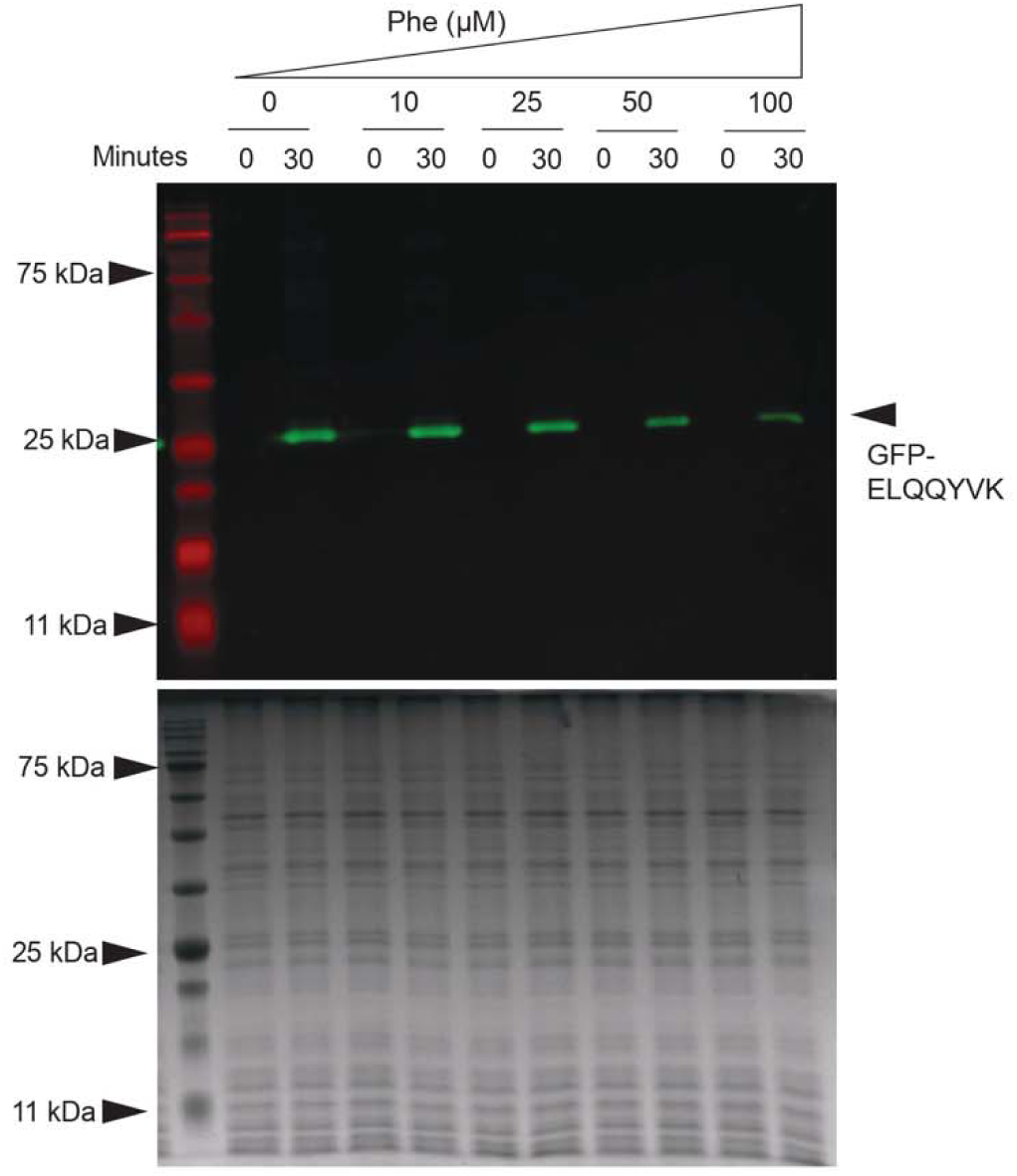
Full immunoblot from Figure 4e, and the Coomassie-stained SDS-PAGE as a loading control.

**Supplementary Figure 15.**
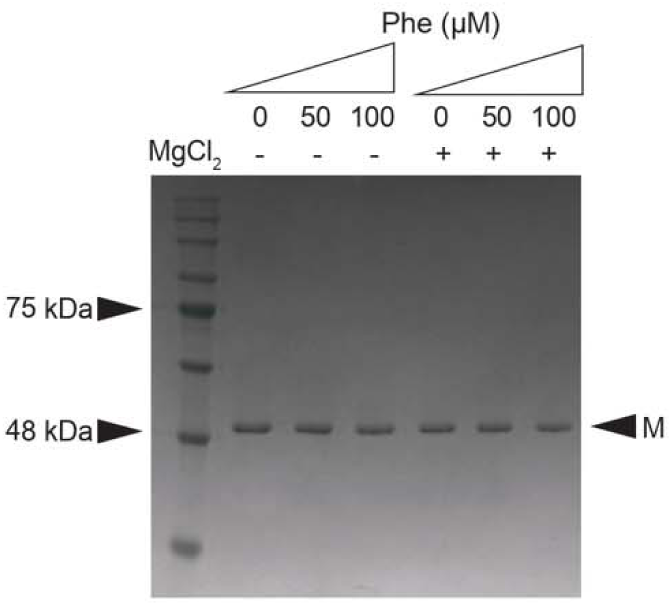
SDS-PAGE of glutaraldehyde-crosslinked ClpX with 0.5 mM ATPγS and 1 mM NaF in the presence or absence of 0.5 mM MgCl_2_ and with increasing phenylalanine. Bands labeled ‘M’ are consistent with monomers.

## Supplementary tables

**Supplementary Table 1.**
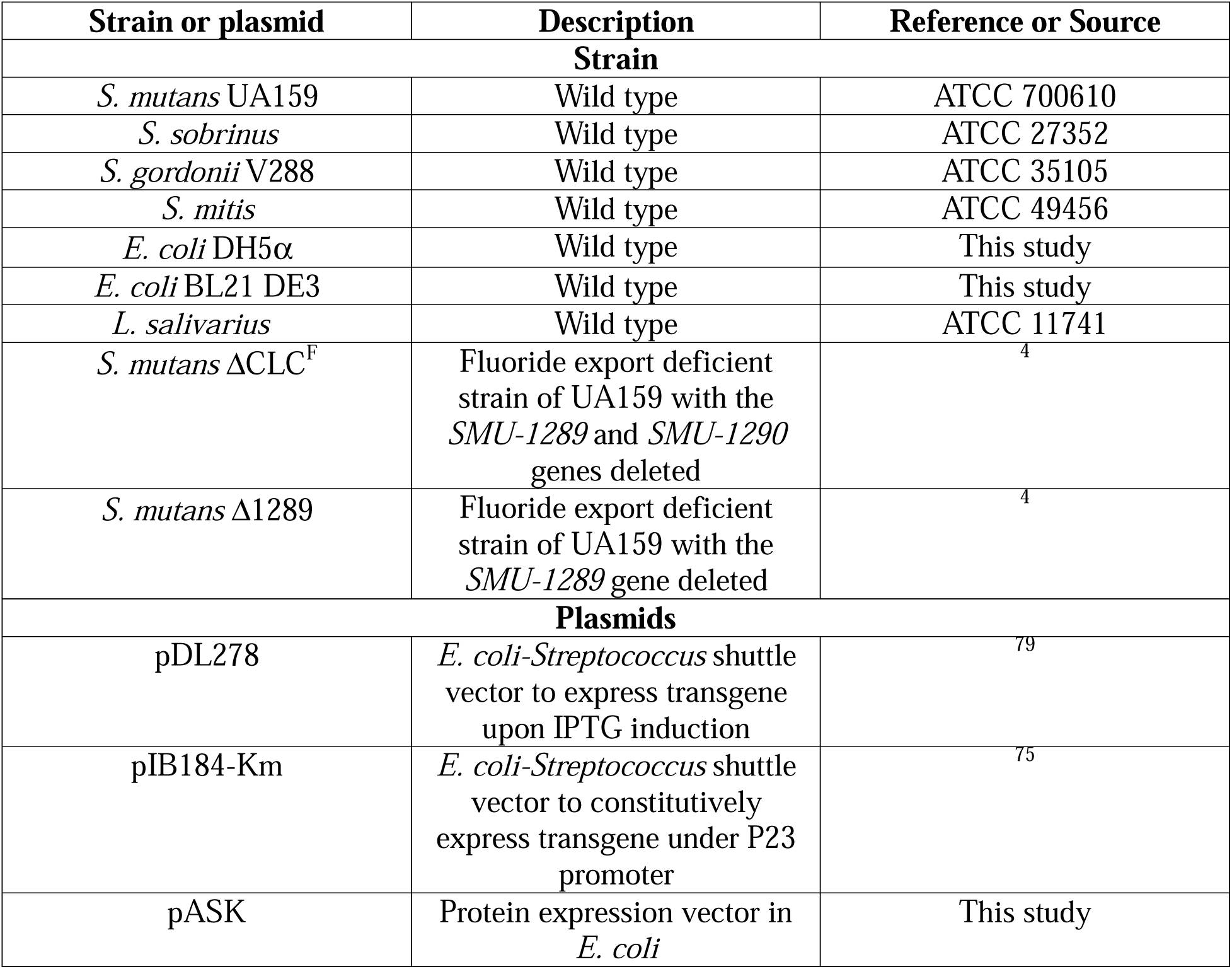
Strains and plasmids used in this study.

| Strain or plasmid | Description | Reference or Source |
| --- | --- | --- |
| **Strain** |  |  |
| *S. mutans* UA159 | Wild type | ATCC 700610 |
| *S. sobrinus* | Wild type | ATCC 27352 |
| *S. gordonii* V288 | Wild type | ATCC 35105 |
| *S. mitis* | Wild type | ATCC 49456 |
| *E. coli* DH5α | Wild type | This study |
| *E. coli* BL21 DE3 | Wild type | This study |
| *L. salivarius* | Wild type | ATCC 11741 |
| *S. mutans* ΔCLC<sup>F</sup> | Fluoride export deficient strain of UA159 with the *SMU-1289* and *SMU-1290* genes deleted | <sup>4</sup> |
| *S. mutans* Δ1289 | Fluoride export deficient strain of UA159 with the *SMU-1289* gene deleted | <sup>4</sup> |
| **Plasmids** |  |  |
| pDL278 | *E. coli-Streptococcus* shuttle vector to express transgene upon IPTG induction | <sup>79</sup> |
| pIB184-Km | *E. coli-Streptococcus* shuttle vector to constitutively express transgene under P23 promoter | <sup>75</sup> |
| pASK | Protein expression vector in *E. coli* | This study |

**Supplementary Table 2.**
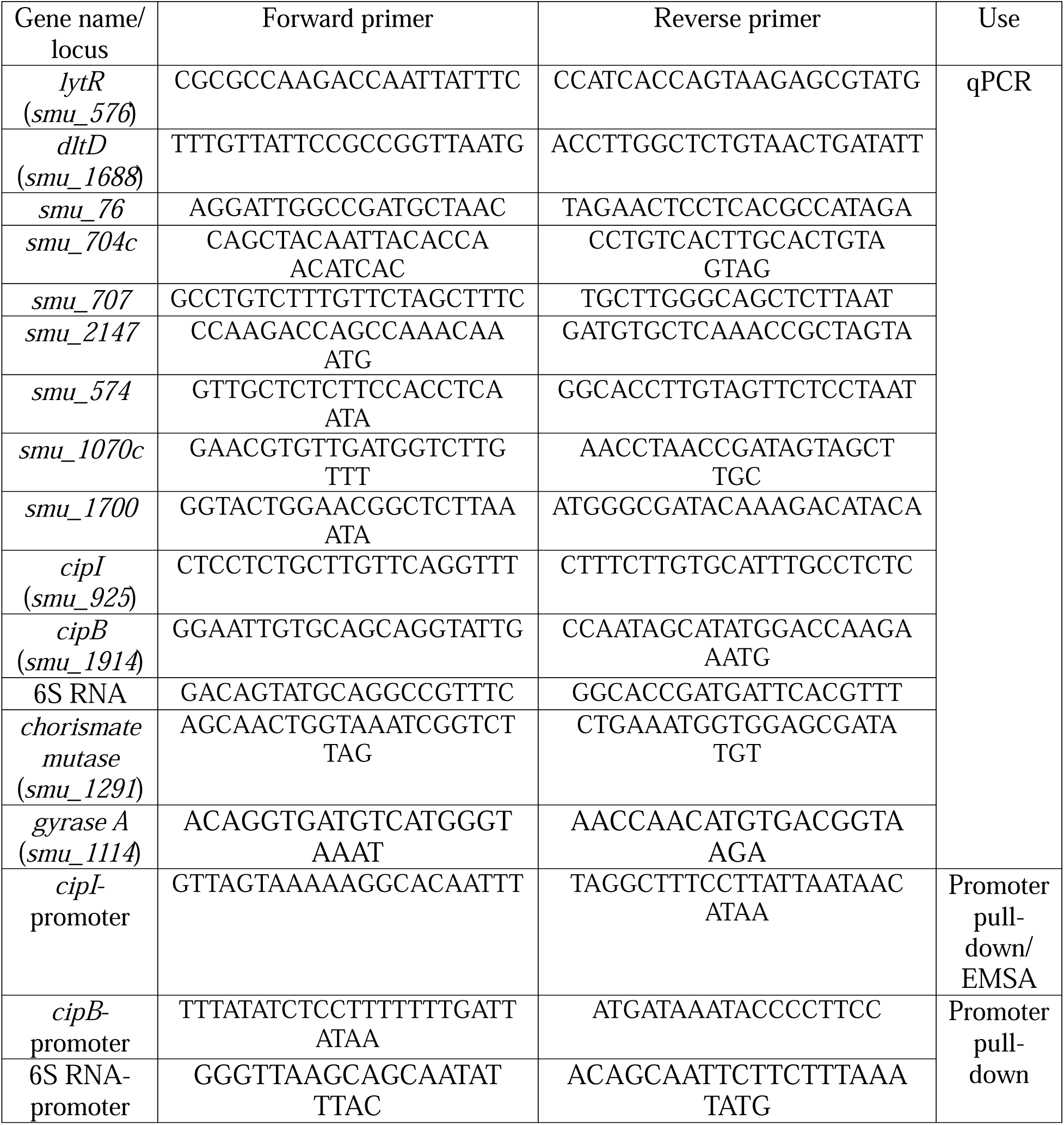
List of primers used in this study.

**Supplementary Table 3.**
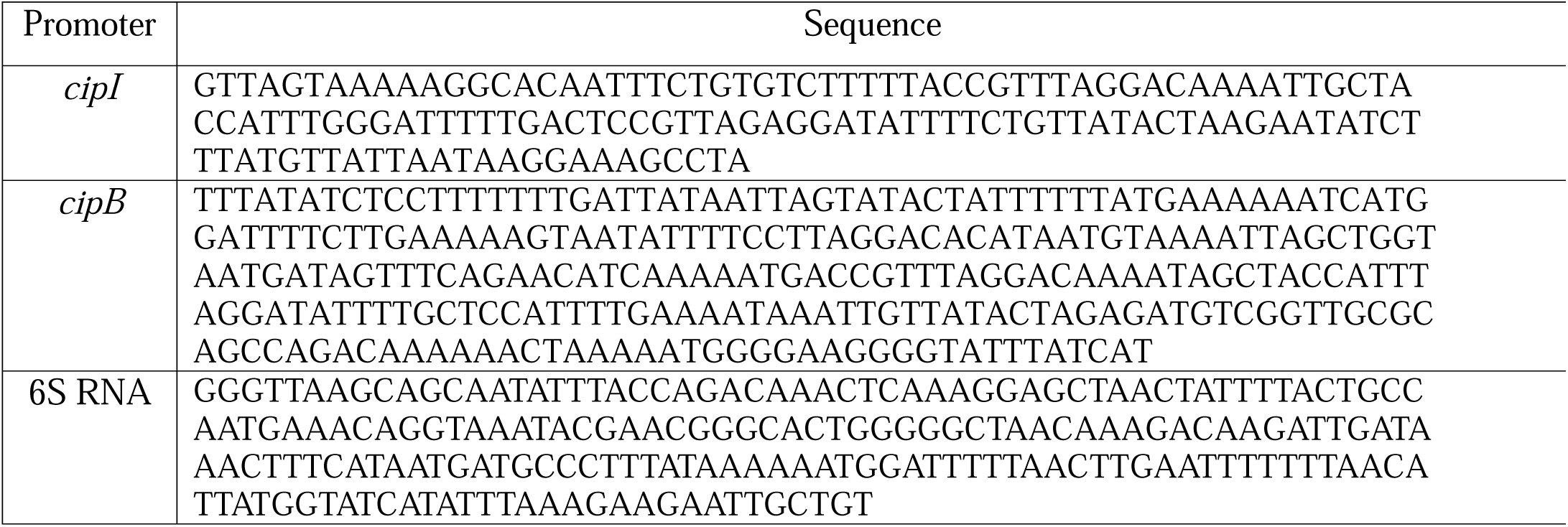
Promoter sequences used for EMSA and pull-down assays.

## Notes

### Competing Interest Statement

The authors have declared no competing interest.

